# Treatment with sodium butyrate induces autophagy resulting in therapeutic benefits for spinocerebellar ataxia type 3

**DOI:** 10.1101/2021.04.30.442119

**Authors:** Maxinne Watchon, Katherine J. Robinson, Luan Luu, Yousun An, Kristy C. Yuan, Stuart K. Plenderleith, Flora Cheng, Emily K. Don, Garth A. Nicholson, Albert Lee, Angela S. Laird

## Abstract

Spinocerebellar ataxia type 3 (SCA3, also known as Machado Joseph disease) is a fatal neurodegenerative disease caused by expansion of the trinucleotide repeat region within the *ATXN3/MJD* gene. Mutation of *ATXN3* causes formation of ataxin-3 protein aggregates, neurodegeneration and motor deficits. Here we investigated the therapeutic potential and mechanistic activity of sodium butyrate (SB), the sodium salt of butyric acid, a metabolite naturally produced by gut microbiota, on cultured SH-SY5Y cells and transgenic zebrafish expressing human ataxin-3 containing 84 glutamine (Q) residues to model SCA3. SCA3 SH-SY5Y cells were found to contain high molecular weight ataxin-3 species and detergent insoluble protein aggregates. Treatment with SB increased activity of the autophagy protein quality control pathway in the SCA3 cells, decreased presence of ataxin-3 aggregates and presence of high molecular weight ataxin-3 in an autophagy-dependent manner. Treatment with SB was also beneficial *in vivo,* improving swimming performance, increasing activity of the autophagy pathway and decreasing presence of insoluble ataxin-3 protein species in the transgenic SCA3 zebrafish. Co-treating the SCA3 zebrafish with SB and chloroquine, an autophagy inhibitor, prevented the beneficial effects of SB on zebrafish swimming, indicating that the improved swimming performance was autophagy-dependent. To understand the mechanism by which SB induces autophagy we performed proteomic analysis of protein lysates from the SB treated and untreated SCA3 SH-SY5Y cells. We found that SB treatment had increased activity of Protein Kinase A and AMPK signalling, with immunoblot analysis confirming that SB treatment had increased levels of AMPK protein and its substrates. Together our findings indicate that treatment with SB can increase activity of the autophagy pathway through a PKA/AMPK-dependent process and that this has beneficial effects *in vitro* and *in vivo*. We propose that treatment with sodium butyrate warrants further investigation as a potential treatment for neurodegenerative diseases underpinned by mechanisms relating to protein aggregation including SCA3.

## Introduction

Spinocerebellar ataxia type 3 (SCA3), also known as Machado-Joseph disease (MJD), is a neurodegenerative disease characterized by a progressive loss of muscle control and movement, leading to wheelchair dependence and decreased lifespan [1]. Clinical symptoms of SCA3 include ataxia, dystonia, rigidity, muscle atrophy and visual and speech disorders [1–3]. SCA3 is the most common of the hereditary ataxias found throughout the world (21-28% of autosomal-dominant ataxia) [4–6], with a high prevalence within the Azores of Portugal [7] and Indigenous communities of north-east Arnhem Land in Australia [8].

SCA3 is caused by inheritance of an expanded CAG repeat region within the *MJD1/ATXN3* gene on chromosome 14 [9–11]. Whilst the *ATXN3* gene of healthy subjects contains a short CAG trinucleotide repeat region (12-40 CAG repeats), this region contains over 40, and as high as 87, CAG repeats in SCA3 patients [7, 9, 12, 13]. However, 44-54 CAG repeats have been considered to be an intermediate length and may not lead to SCA3 [14]. The *ATXN3* gene encodes the ataxin-3 protein, with the CAG repeat region encoding a polyglutamine (polyQ) tract towards the C-terminus of the protein [12, 13]. Neuropathological staining of patient brain samples often reveals the presence of neuronal intranuclear inclusions (NII) containing the ataxin-3 protein [2, 15, 16] and extraction of these proteins has revealed the presence of full-length ataxin-3 protein, as well as smaller ataxin-3 protein fragments [17].

Whilst the function of the ataxin-3 protein is not fully understood it is known to function as a deubiquitinating (DUB) enzyme [1] and can affect levels of transcription through interactions with co-activators and histones [18–20]. It has been previously reported that ataxin-3 is a histone binding protein and that polyQ expansion within the protein increases the extent of that binding, in turn affecting histone acetylation [18]. Furthermore, it has also been suggested that mutated polyQ proteins can also inhibit the function of histone acetyltransferases [21–23]. Indeed, cell culture and animal models, such as *Drosophila*, transgenic mice and recently our transgenic zebrafish expressing polyQ expanded human ataxin-3, have been reported to have histone hypoacetylation [22, 24–26]. However, this histone hypoacetylation has not been reported to be present in all models of the disease [21].

One therapeutic strategy that has been explored for SCA3 is treatment with drugs that may increase levels of histone acetylation, called histone deacetylase (HDAC) inhibitors. HDACs are a class of enzymes that remove acetyl groups from ε-N-acetyl lysine located on histones, causing histones to bind DNA more tightly and prevent transcription [27, 28]. Previous studies have reported that treatment with HDAC inhibitors can have neuroprotective effects for the treatment of SCA3 [21, 22, 24–26, 29] and other neurodegenerative diseases including Huntington’s disease and Alzheimer’s disease [30–33]. Compounds with this HDAC inhibitor capacity include suberoylanilide hydroxamic acid (SAHA), trichostatin A, resveratrol, valproic acid and sodium butyrate [27, 34–38]. Moreover, a phase I/II clinical trial was conducted for sodium valproate in SCA3 patients with minimal adverse events [39].

Sodium butyrate (SB) is a sodium salt of butyric acid, a short-chain fatty acid that is produced within the gut by intestinal microbiota during fermentation of undigested dietary carbohydrates and fibre [28, 40]. Whilst butyric acid is critical for intestinal homeostasis, it also exerts a wider range of effects including inhibition of cell proliferation, downregulation of pro-inflammatory effectors, repression of gene expression and HDAC inhibition, resulting in increased acetylation of histones 3 and 4 [25, 40]. Recently, a combined treatment of taurursodiol and phenylbutyrate, another derivative of butyric acid with HDAC inhibitor effects, has been found to improve survival in a double-blind, placebo controlled clinical trial of amyotrophic lateral sclerosis (ALS) patients [41]. Since then, this combination therapy has received FDA approval for the treatment of ALS. As butyrate can elicit such a wide spectrum of positive effects, it is likely butyrate has multiple distinct mechanisms of action [40, 42], many of which are yet to be fully elucidated.

In this study, we aimed to determine the efficacy of SB in alleviating SCA3 pathology in cell culture and zebrafish models of SCA3. Furthermore, we aimed to elucidate the underlying mechanism by which SB rescues SCA3 disease phenotypes. Enhancing our understanding of the mechanisms of action underlying the neuroprotective effects induced by SB could provide therapeutic implications for a wide range of neurodegenerative and proteinopathy diseases. We hypothesized that, in addition to improving transcriptional dysregulation, SB can ameliorate SCA3 phenotypes via induction of the autophagy protein quality control pathway. Increased activation of the autophagy pathway may act to degrade or remove pathological ataxin-3 oligomeric species or aggregates in cell cultures, alleviating neurotoxicity. Importantly, here we demonstrate that administration of SB can induce autophagy and decrease SCA3 disease phenotypes both *in vitro* and *in vivo*, highlighting the therapeutic potential of autophagy inducers for the treatment of SCA3 and other related neurodegenerative diseases.

## Results

### A SCA3 SH-SY5Y model recapitulates phenotypes found in human SCA3 patients, including presence of protein aggregates and histone hypoacetylation

Immunoblot analysis of neuron-like (neuroblastoma, SH-SY5Y) cells stably expressing human ataxin-3 or an empty vector control revealed the expression of endogenous ataxin-3, ataxin-3-28Q and ataxin-3-84Q at appropriate sizes (45 kDa, 50 kDa and 65 kDa respectively, Figure. 1A) (Uncropped images found in Supp. Figure. 1). Expression of ataxin-3 monomers did not differ between ataxin-3 28Q and ataxin-3 84Q cells, but as expected, was greater than in the empty vector control cells (Figure. 1B). Cells expressing ataxin-3-84Q were found to contain an additional high molecular weight (HMW) ataxin-3 band, at approximately 140 kDa, which was not present in cells expressing ataxin-3-28Q (Figure. 1C). We also found that these HMW bands were not present if we instead transfected cells with alternative constructs to express human ataxin-3 (data not shown).

**Figure 1.**
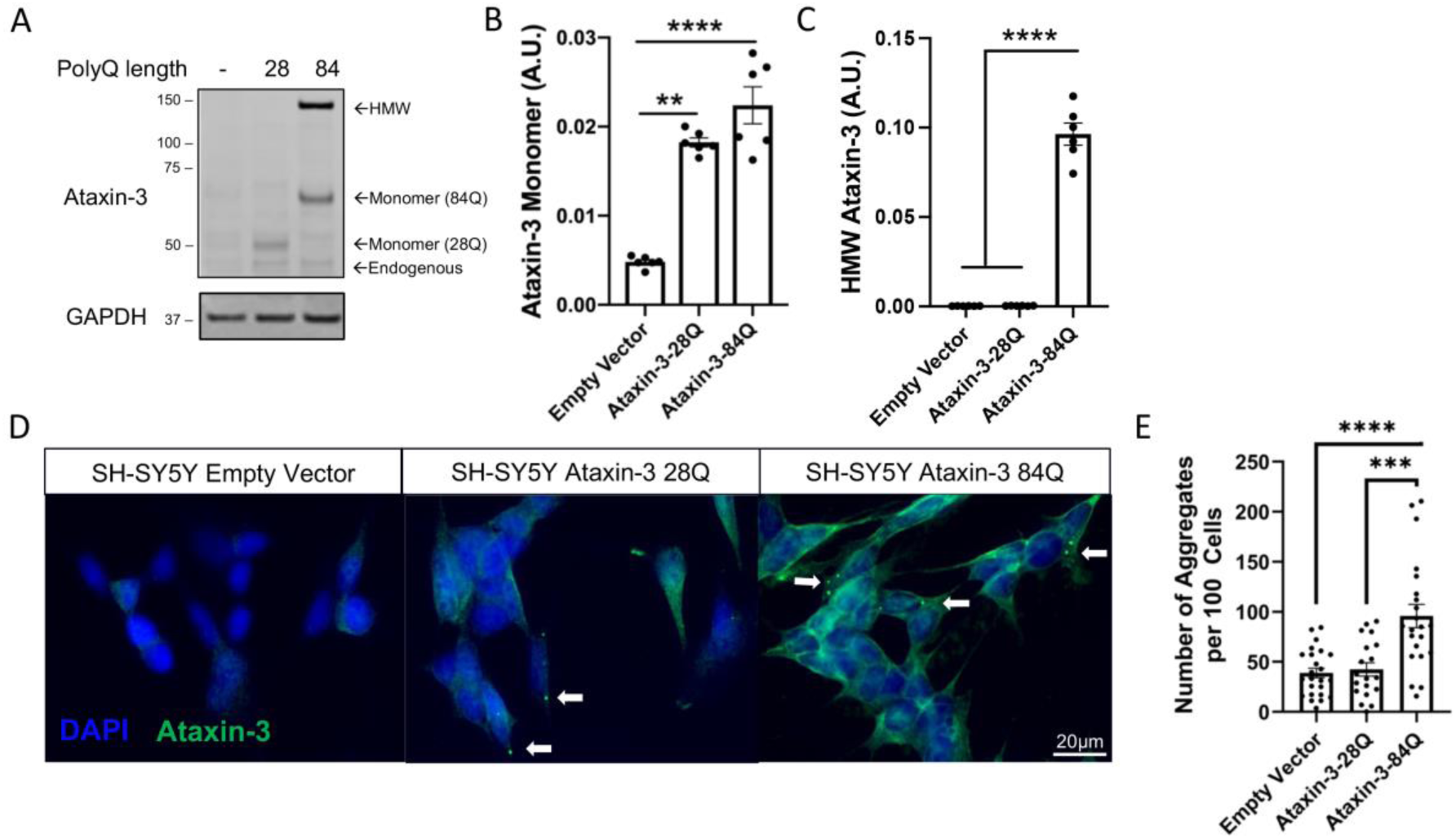
SH-SY5Y cell culture modelling revealed the presence of ataxin-3 aggregates. A) Immunoblotting SH-SY5Y cells with ataxin-3 confirmed expression of human ataxin-3 28Q and 84Q as well as endogenous ataxin-3. An additional high molecular weight (HMW) band above the 84Q monomeric band was found. (B) Quantification of human ataxin-3 monomers indicated increased ataxin-3 levels in ataxin-3 28Q and 84Q compared to the vector control (p < 0.001 and p = 0.0013 respectively; n=6), and (C) quantification of the HMW ataxin-3 bands revealed that more HMW ataxin-3 was present in the ataxin-3 84Q cells compared to the other genotypes (p < 0.0001; n = 6). (D) Stably transfected SH-SY5Y cells were confirmed to express human ataxin-3 after immunostaining for ataxin-3. Cells expressing an empty vector control did not display aggregated ataxin-3, however cells expressing either human ataxin 3 28Q, or ataxin-3 84Q displayed ataxin-3 positive aggregates (indicated by white arrows, scale bar: 20 µm). (E) Quantification of the number of aggregates present revealed that ataxin-3 84Q cells possessed a higher number of aggregates compared to ataxin-3 28Q and the vector control (p = 0.001 and p < 0.0001 respectively, n = 19-24). Results are shown as mean ± SEM. Statistical analysis was performed using a one-way ANOVA followed by a Tukey’s post hocc test.

We confirmed that this HMW band contained ataxin-3 by comparing the location of the immunoreactive band of HMW ataxin-3 on the immunoblot with a Coomassie stained 1D SDS PAGE gel that was run in parallel to the immunoblot. This region was excised, in-gel trypsin digestion was performed, and tryptic peptides were analyzed by LC-MS/MS. We confirmed that this immunoreactive band indeed contained ataxin-3 (Supp. Figure. 2, Supp. Table 1).

Immunostaining of the SH-SY5Y cells stably expressing human ataxin-3, or an empty vector control, showed expression of ataxin-3 throughout the cytoplasm and nucleus of the cell (Figure. 1D). Previously, we demonstrated that SH-SY5Y cells transiently transfected with EGFP-fused human ataxin-3-84Q carry Triton-X insoluble protein aggregates that can be detected by a flow cytometry protocol called FloIT [43]. Upon immunostaining the stably expressing ataxin-3 cells for ataxin-3, the presence of cytoplasmic ataxin-3-positive puncta was revealed in cells expressing ataxin-3-28Q and −84Q (Figure. 1D; white arrows). Automated quantification of the number of potential ataxin-3 protein aggregates (puncta with a diameter of 2.25-6 µm) within the images revealed that cells stably expressing ataxin-3-84Q had approximately double the number of puncta than cells expressing ataxin-3-28Q or an empty vector control (Figure. 1E).

We have previously demonstrated that our transgenic SCA3 zebrafish exhibited alterations in acetylation of histone 3 at lysine 9 (ac-H3K9) and histone 4 at lysine 5 (ac-H4K5) compared to a non-transgenic control zebrafish [26]. In this study, we sought to confirm whether this finding is consistent in the SH-SY5Y cells expressing polyQ expanded human ataxin-3. Interestingly, we found that SH-SY5Y cells stably expressing ataxin-3-84Q had decreased ac-H3K9 compared to the ataxin-3 28Q expressing cells, whilst similar levels of ac-H4K5 were detected across examined genotypes (Supp. Figure. 3).

### Treatment with sodium butyrate increases histone acetylation and decreases ataxin-3 aggregates in SCA3 cell culture models

To investigate the therapeutic potential of sodium butyrate (SB), we treated SH-SY5Y cells stably expressing human ataxin-3 84Q with SB (3 mM) for 72 hours and analysed whether ataxin-3-84Q expressing cells displayed increased histone acetylation following SB treatment. Immunoblotting for the ac-H3K9 and ac-H4K5 revealed that SB treatment produced a 50- and 7-fold increase in both ac-H3K9 and ac-H4K5, respectively (Figure. 2A-C; Uncropped images found in Supp. Figure 4). Treatment with SB for 3 days was also found to decrease the number of ataxin-3 aggregates detected by ataxin-3 immunostaining by approximately 30 %, when compared to vehicle treated ataxin-3 84Q expressing cells (Figure. 2D, E). Immunoblot analysis revealed that SB treatment also decreased the presence of the HMW ataxin-3 species by approximately 25 % (Figure. 2F, G) and produced a 1.5-fold increase in the amount of monomeric ataxin-3 species of present (Figure. 2H), suggesting the induction of a protein quality control pathway that is targeting HMW ataxin-3 (Uncropped images found in Supp. Figure. 4). Interestingly, flow cytometric analysis of aggregates (FloIT) revealed an increase in the number of detergent insoluble GFP^+^ particles with only 24 hours of treatment (Supp. Figure. 5A-B).

**Figure 2:**
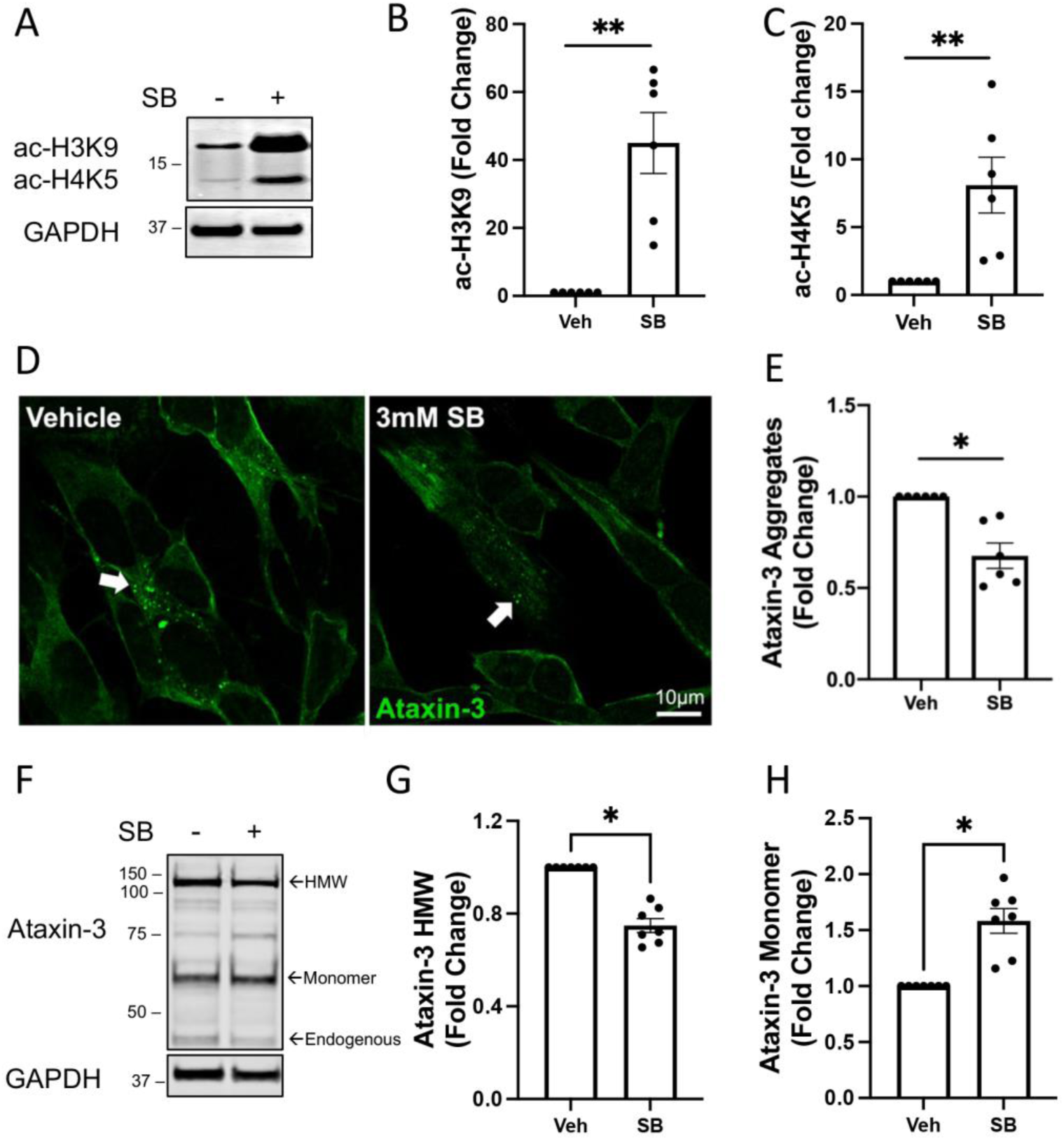
Treatment with sodium butyrate increases histone acetylation and affects ataxin-3 aggregation in SCA3 SH-SY5Y cells. (A) Western blots of SH-SY5Y stably expressing ataxin-3 84Q cells treated with 3 mM SB or vehicle control were probed for acetylated histone 3 at lysine 9 (ac-H3K9) and acetylated histone 4 at lysine 5 (ac-H4K5). (B) Densitometry of ac-H3K9 showed an increase with SB treatment compared to vehicle treatment (p = 0.0312). (C) Densitometry of ac-H4K5 revealed an increase with SB treatment compared to vehicle control (p = 0.0022, n=6). (D) Immunostaining for ataxin-3 revealed a reduction in the amount of ataxin-3 positive aggregates after treatment with SB, compared to vehicle treatment (white arrows, scale bar: 10 µm). (E) Quantification of the number of ataxin-3 positive aggregates revealed decreased aggregates in SB-treated ataxin-3 84Q cells compared to the vehicle treated control (p = 0.0022, n=6). (F) Western blot of SH-SY5Y stably expressing ataxin-3 84Q cells treated with 3 mM SB or vehicle were probed for ataxin-3. (G) Densitometry of high molecular weight (HMW) ataxin-3 indicated that decreased HMW ataxin-3 was present following treatment with SB compared to vehicle control (p = 0.0156, n = 7). (H) Quantification of the ataxin-3 monomer levels following SB treatment increased compared to the vehicle control (p = 0.0156, n=7). Data represents mean ± SEM. The statistical analysis was performed by Wilcoxon test.

### Treatment with sodium butyrate increases histone acetylation and improves swimming of SCA3 zebrafish

Previously, we developed a transgenic zebrafish model of SCA3 with the intent to screen drug candidates for the treatment of SCA3 [44]. From this model we found that our SCA3 zebrafish larvae expressing EGFP-tagged human ataxin-3 84Q had altered levels of histone acetylation, namely ac-H3K9 and ac-H4K5, compared to non-transgenic zebrafish [26]. Since SB is a known HDAC inhibitor, EGFP-ataxin-3-84Q zebrafish were treated with SB (250 µM, 500 µM and 1 mM) from 24 hours post fertilization (hpf) to 6 days post fertilisation (dpf) to determine whether levels of histone acetylation could be increased. Immunoblot analysis confirmed that 6 dpf transgenic zebrafish expressed human ataxin-3 protein of appropriate sizes (72 kDa and 84 kDa for EGFP-ataxin-3 23Q and 84Q respectively), as well as endogenous zebrafish ataxin-3 (34 kDa; Figure. 3A; Uncropped images found in Supp. Figure. 6). Lower molecular weight ataxin-3-positive cleavage fragments were also present, consistent with our previous findings [44]. Further, immunoblotting for ac-H3K9 and ac-H4K5 revealed a dose dependent increase in levels of acetylated of histone 3 and histone 4 with SB treatment (Figure. 3A). This was confirmed when quantifying the amount of ac-H3K9 following treatment with 500 μM and 1 mM SB (Figure. 3B) and ac-H4K5 following 1 mM SB (Figure. 3C), compared to the vehicle-treated control. Quantification of levels of ataxin-3 revealed that there was also a 3-fold and 1.6-fold increase in the presence of full-length (monomeric) and cleaved human ataxin-3 after treatment with both 500 µM and 1 mM SB (Figure. 3D, E).

**Figure 3.**
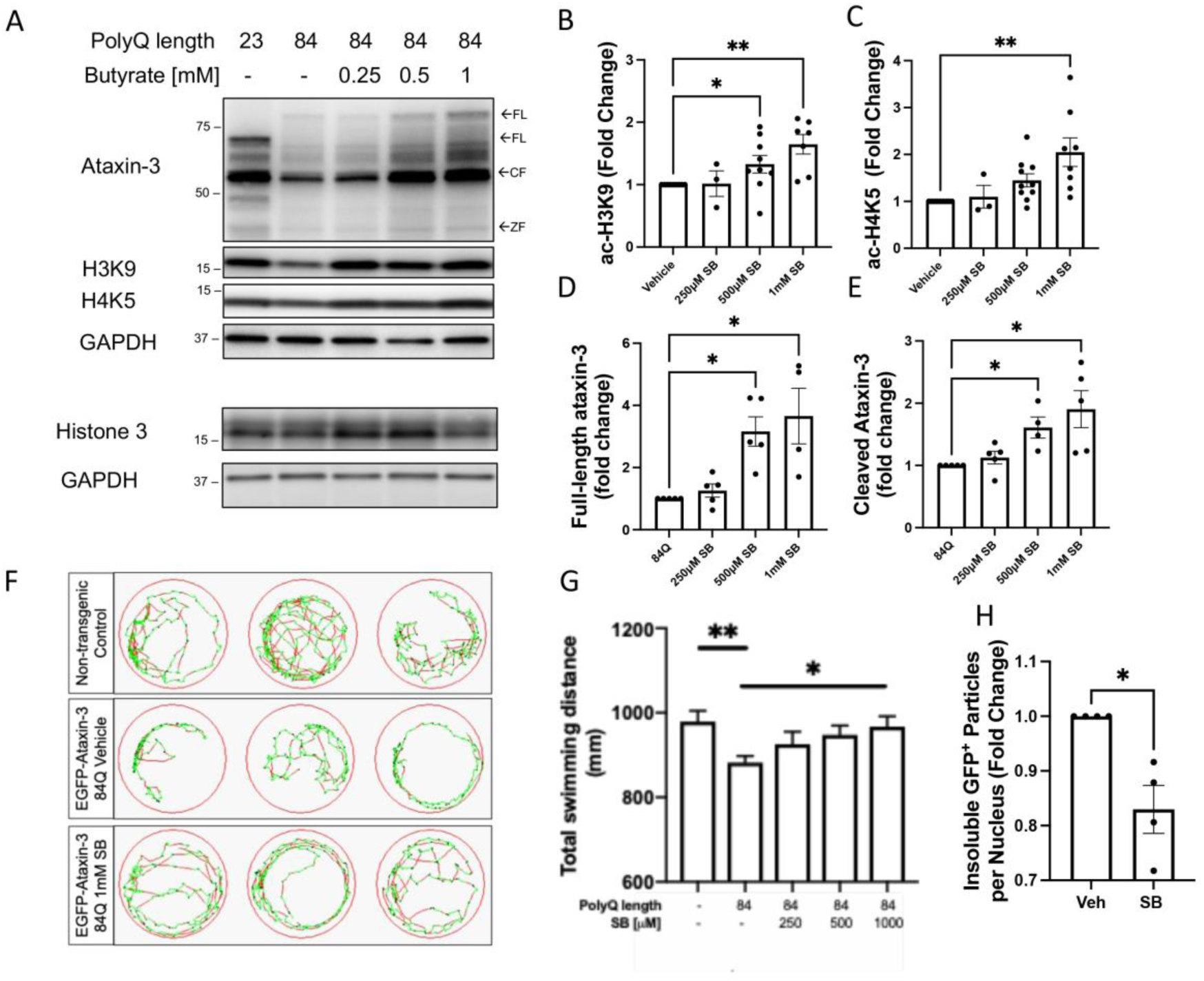
Sodium butyrate treatment increases histone acetylation and rescues the motor phenotype in SCA3 zebrafish. Zebrafish expressing ataxin-3 84Q were treated with 250 µM, 500 µM or 1 mM sodium butyrate (SB), or vehicle control, between 1-6 days post fertilization (dpf). (A) Protein lysates of the 6 dpf zebrafish treated either SB or the vehicle control underwent western blotting and checked for ataxin-3, acetylated histone 3 at lysine 9 (ac-H3K9) and acetylated histone 4 at lysine 5 (ac-H4K5). Histone 3 was probed as a control. (B) Quantification of ac-H3K9 revealed a dose-dependent increase following SB treatment (500 µM, p = 0.0481; 1 mM, p = 0.0014, n=3-9), whilst (C) quantification of ac-H4K5 revealed 1 mM SB was able to increase histone acetylation compared to the vehicle control (p = 0.0018, n = 3-10). (D) Quantification of full-length (FL) ataxin-3 revealed an increase with 500 µM and 1 mM SB (p = 0.039 and p = 0.0452 respectively, n = 4-5) whilst (E) levels of the cleavage fragment of ataxin-3 (CF) also demonstrated an increase with 500 µM and 1 mM SB treatment (p = 0.0452 and p = 0.0109 respectively, n = 4-5). (F) Swimming trajectories of 6 dpf zebrafish after treatment with vehicle versus 1 mM SB treatment (Green, slow movement; red, fast movement). (G) Motor behaviour analysis showed vehicle treated ataxin-3 84Q zebrafish swam shorter distances compared to the non-transgenic controls (p = 0.0085) whilst 6 dpf ataxin-3-84Q larvae treated with 1 mM SB ameliorated the motor dysfunction (p = 0.0393, n = 64-158). (H) Flow cytometric analysis of the insoluble GFP^+^ particles revealed a decrease with SB treatment compared with vehicle (p = 0.03, n = 4 group replicates). Data represents mean ± SEM. Statistical analysis was performed by a non-parametric one-way ANOVA (Kruskal Wallis) test and comparison between vehicle versus SB was analysed using an unpaired student t-test.

Next, we examined whether SB treatment could prevent motor impairment displayed in mutant ataxin-3 zebrafish at 6 dpf. We first examined whether treatment with these concentrations of SB would affect survival and behaviour of non-transgenic zebrafish at 6 dpf. SB treatment, irrespective of concentration, did not affect the survival or behaviour of the non-transgenic zebrafish (Supp. Figure 7). Therefore, upon examination of the movement of the SCA3 zebrafish during the escape response to darkness motor assay revealed, as reported previously [44], that zebrafish expressing ataxin-3 with a polyglutamine expansion swam shorter distances than non-transgenic controls (Figure. 3F, G). Treatment with SB produced a dose-dependent increase of swimming distance, with 1 mM SB producing a significant rescue of the swimming of the EGFP-ataxin-3 84Q zebrafish (Figure. 3G).

Flow cytometric analysis was also used to confirm whether SB treatment affected the number of insoluble EGFP-ataxin-3 aggregates. Previously, we have been able to show that zebrafish larvae expressing EGFP-ataxin-3-84Q develop protein aggregates detectable via whole mount confocal imaging and a modified FloIT protocol, as early as 2 dpf [43]. SB treatment from 1 to 6 dpf resulted in a 17 % decrease in the number of insoluble EGFP-ataxin-3 particles compared to vehicle treated controls (Figure. 3H). In contrast, no decrease in insoluble EGFP-ataxin-3 particles was found after short-term SB treatment (from 1 - 2 dpf) (Supp. Figure. 5C, D).

### Autophagy pathway is impaired in SCA3 SH-SY5Y cells and sodium butyrate treatment induces autophagy and aids removal of high molecular weight ataxin-3

As we detected ataxin-3 aggregates within the cells expressing ataxin-3 84Q, and SB treatment was able to reduce the number of these ataxin-3 aggregates and high molecular weight (HMW) ataxin-3 bands, we next explored whether the protein quality control pathway, macroautophagy (further known as autophagy) was activated in these cell cultures. It has been shown previously that autophagy pathway substrates can be sequestered into ataxin-3 protein aggregates and autophagic flux can be dysfunctional, in SCA3 patient brain samples and patient fibroblasts, respectively [45–48]. We reported previously that the SH-SY5Y cells stably expressing polyQ expanded ataxin-3 have impaired autophagy dynamics [49]. To examine whether the reduction of HMW ataxin-3 species was mediated by SB-induced autophagy, SH-SY5Y ataxin-3 84Q cells were co-treated with SB (3 mM) and an autophagy inhibitor, 3-Methyladenine (3MA, 5 mM), for 72 hours (Figure. 4A; Uncropped images found in Supp. Figure 8). Whilst SB treatment decreased the amount of HMW ataxin-3 present compared to vehicle control by approximately 25 %, co-treatment with SB and 3MA (SB+3MA) together prevented the decrease in the amount of HMW ataxin-3 compared to vehicle-treated cells (Figure. 4A, B).

**Figure 4.**
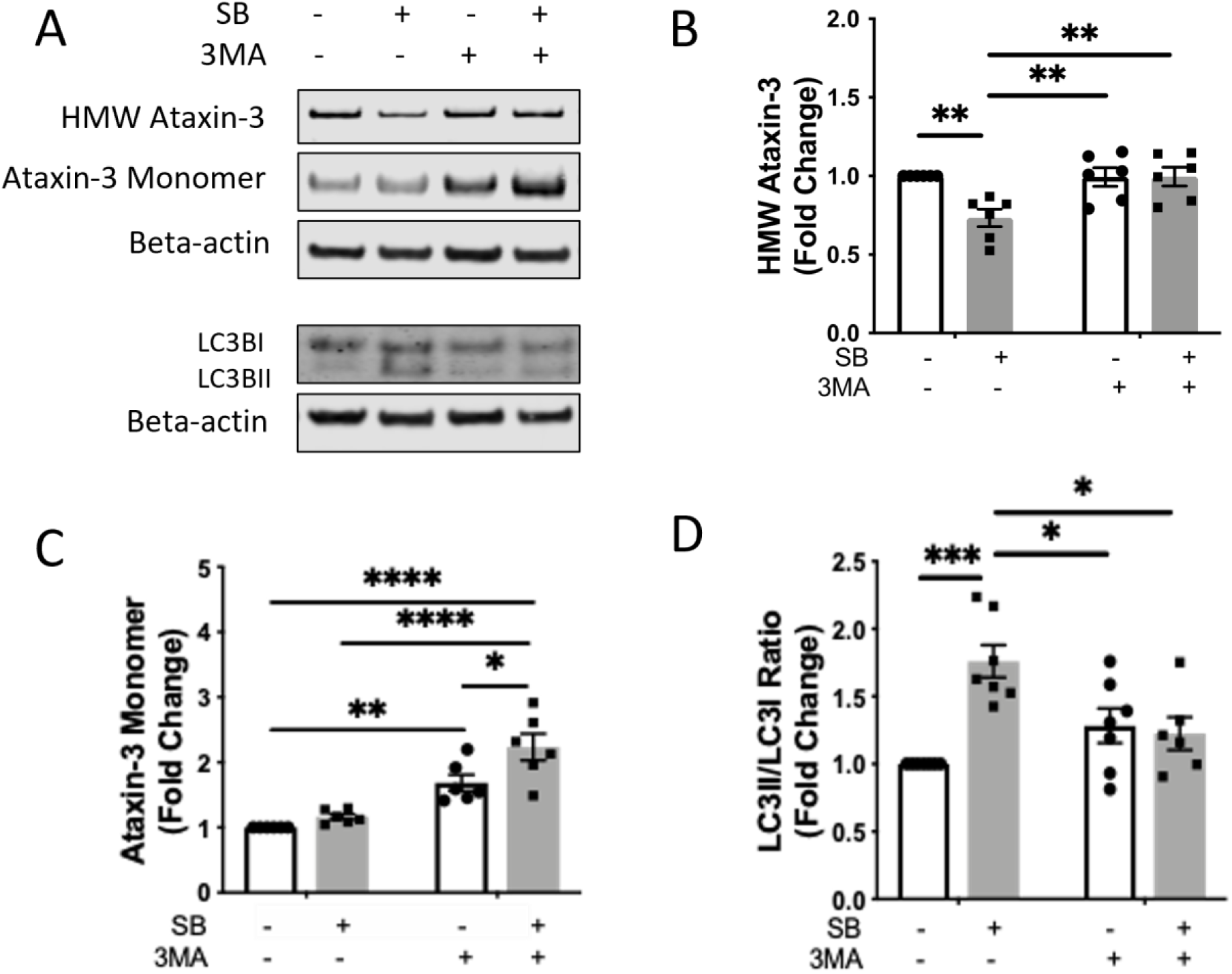
Removal of high molecular weight (HMW) ataxin-3 by sodium butyrate is mediated by autophagy. (A) Western blots were performed on protein lysates of SH-SY5Y cells stably expressing ataxin-3 84Q treated with sodium butyrate (SB, 3 mM) and/or 3MA (5 mM) for 72 hrs and probed for ataxin-3 and LC3B. (B) Quantification of HMW ataxin-3 indicated that SB-treatment reduced the amount of HMW ataxin-3, compared to vehicle treatment (p = 0.0070), and addition of 3MA treatment alone or as a co-treatment prevented the removal of the HMW ataxin-3 (p = 0.0086 and p = 0.0079 respectively, n=6). (C) Quantification of the ataxin-3 monomeric species increased with 3MA treatment compared to the vehicle control (p = 0.0036). SB+3MA co-treatment further increased ataxin-3 levels compared to vehicle, SB treatment alone and 3MA treatment alone (p < 0.0001, p < 0.0001 and p = 0.0214 respectively, n=6). (D) Quantification of the LC3II/LC3I ratio revealed an increased ratio with SB treatment (p = 0.0002) and the addition of 3MA alone or SB+3MA co-treatment prevented this decrease in LC3II/LC3I (p = 0.0176 and p = 0.0098 respectively; n=6-7). Data represents mean ± SEM. Statistical analysis performed was a two-way ANOVA followed by a Tukey post-hoc analysis.

Levels of monomeric ataxin-3 increased with 3MA treatment alone, as well as SB+3MA co-treatment by ≥1.7 fold (Figure. 4C). Together, these findings indicate that 3MA prevented the capacity of SB to decrease HMW ataxin-3, and that the removal of HMW ataxin-3 was dependent on SB-mediated autophagic activity.

To examine whether SB treatment was increasing autophagic flux, immunoblots containing protein lysates of the SH-SY5Y cells expressing ataxin-3 84Q treated with either vehicle or SB and/or 3MA were probed with a marker of autophagosomes, LC3B (Figure. 4A). Cells treated with SB displayed a 1.7-fold increase in the LC3II/LC3I ratio compared to vehicle controls (Figure. 4D). In comparison, co-treatment with SB+3MA prevented this increase, indicating that it was dependent on the autophagy pathway (Figure. 4D). In a similar manner, we found that co-treatment with SB and another autophagy inhibitor, bafilomycin (Baf A1), which acts to prevent the degradation of autophagosomes, resulted in an increase in LC3II/LC3I ratio compared to following Baf A1 treatment alone, confirming that SB-treatment increases autophagosome formation (Supp. Figure 9).

### Sodium butyrate treatment induces the autophagy pathway in the transgenic SCA3 zebrafish and increases zebrafish swimming in an autophagy-dependent manner

To confirm that autophagy was also being induced by SB treatment *in vivo,* the transgenic SCA3 zebrafish were treated from 1-6 dpf with either vehicle, SB (1 mM), an autophagy inhibitor (chloroquine, 1.5 mM) or co-treated with SB and chloroquine. Protein lysates of groups of larvae from these treatment groups were then subjected to western blot analysis and probed for LC3B (Figure. 5A; Uncropped images found in Supp. Figure 10).

**Figure 5:**
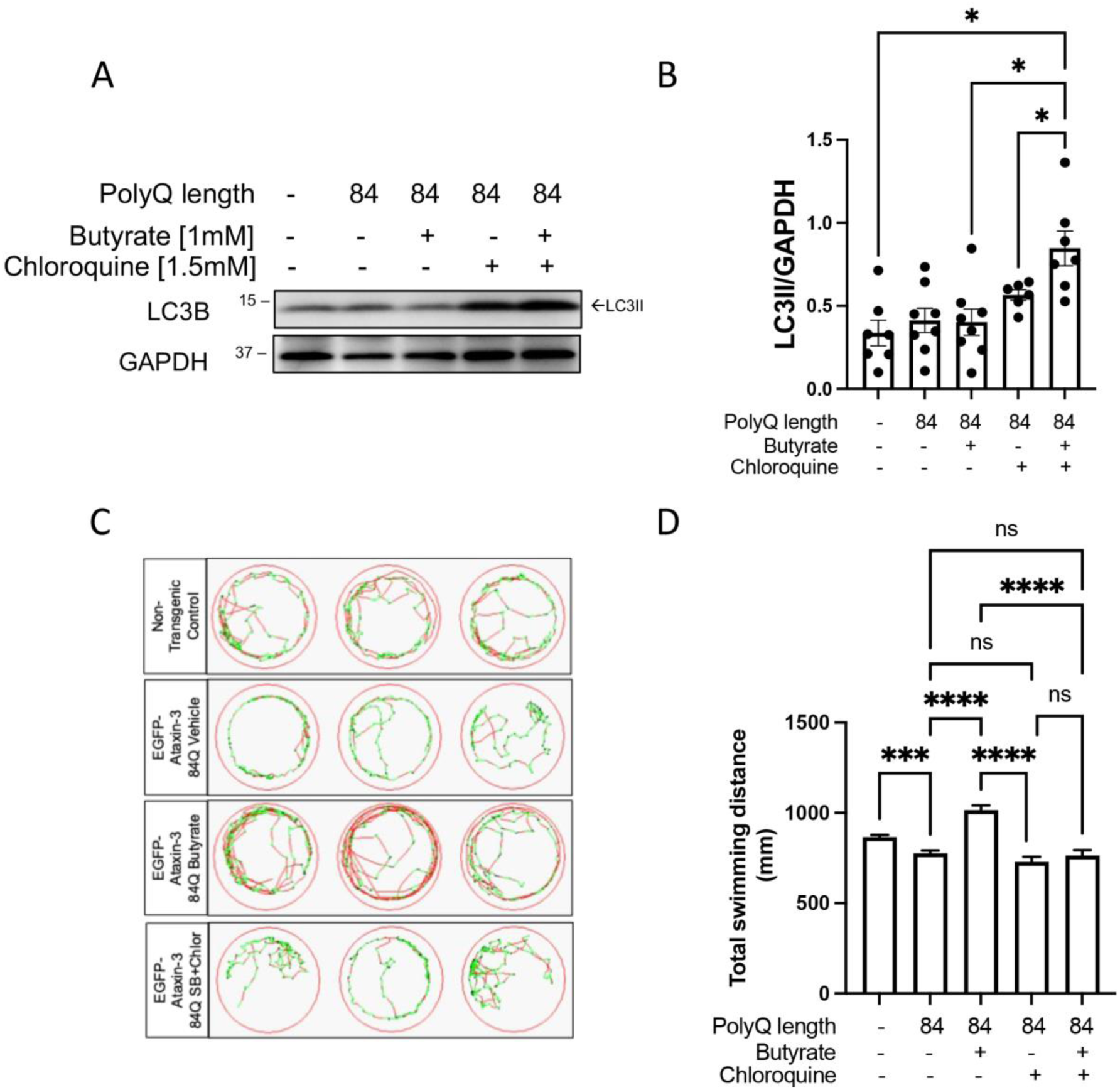
Treatment with sodium butyrate induces autophagy in SCA3 zebrafish model producing beneficial effects on zebrafish swimming. (A) Western blots were performed on protein lysates from non-transgenic controls and ataxin-3 84Q zebrafish larvae treated with either 1 mM sodium butyrate (SB) alone, 1.5 mM chloroquine alone or co-treatment of SB and chloroquine, between 1-6 days post fertilization (dpf). The immunoblots were probed for LC3B. (B) Quantification of LC3II levels after treatment with 1mM SB and/or 1.5 mM chloroquine found an increase with SB and chloroquine treatment compared to chloroquine treatment alone, SB treatment alone and the non-transgenic controls (p = 0.0475, p = 0.0395 and p = 0.0303 respectively; n=6-8). (C) Swimming trajectories of non-transgenic and SCA3 zebrafish treated with either vehicle control, 1 mM SB, 1.5 mM chloroquine or SB/1.5 mM chloroquine and show that SB increases the amount of fast swimming and SB/chloroquine co-treatment decreases this (green, slow movement; red, fast movement). (D) Quantification of the total distance swum by each animal demonstrates that vehicle treated SCA3 zebrafish has decreased swimming distance (p = 0.0009), SB treatment rescues the motor impairment seen in vehicle treated ataxin-3 84Q zebrafish (p< 0.0001) and those receiving 1.5 mM chlor or SB/chlor co-treatment (p < 0.0001 and p < 0.0001 respectively, n=81-260). Data represents mean ± SEM. Statistical analysis was performed using a one-way ANOVA followed by a Tukey post-hoc test. ns – non-significant

Densiometric analysis revealed that SB treatment alone produced similar levels of LC3II to vehicle treated larvae (Figure. 5B). Levels of LC3II increased in co-treatment of SB and chloroquine compared to the non-transgenic control, SB treatment alone and chloroquine treatment alone. The finding of increased LC3II levels following co-treatment with SB and chloroquine, when compared to chloroquine alone, indicates that autophagy is being induced upstream of the autophagy blockade produced by chloroquine. Taken together, these findings confirm that autophagy was induced in the SCA3 zebrafish by the SB treatment.

To confirm that the improved swimming phenotype of the SCA3 zebrafish produced by SB treatment was due to the induction of autophagy, we performed a co-treatment experiment followed by motor behavior analysis. The SCA3 zebrafish larvae were treated with SB (1 mM), chloroquine (1.5 mM), or both, from 1-6 dpf, prior to motor tracking. Vehicle treated Ataxin-3 84Q zebrafish had a shorter swimming distance compared to the non-transgenic control and treatment with SB alone again resulted in an increase in the distances swum by the SCA3 zebrafish, compared to vehicle treatment. The addition of 1.5 mM chloroquine with SB treatment prevented this improvement in motor function (Figure. 5C, D). This indicates that chloroquine treatment was able to suppress the beneficial effect of SB, and therefore that the beneficial effect of SB treatment was indeed autophagy dependent.

### Sodium butyrate mediated autophagy induction is predicted to occur through activation of protein kinase A and AMPK pathways

To investigate the potential mechanisms by which SB was inducing autophagy, we performed proteomic analysis to compare lysates extracted from SH-SY5Y cells expressing ataxin-3 84Q, treated for 72 hours with either SB (3 mM) or vehicle control. From triplicate analysis, we identified 5143 and 5110 proteins from vehicle and treated cells respectively, with 4966 (93.9 %) proteins identified in common. We therefore identified 177 (3.3 %) and 144 (2.7 %) unique proteins between the vehicle and SB treated cell lysates. There did not appear to be obvious differences in the protein classes that were identified between the vehicle and treated samples. Therefore, we carried out label-free quantitative proteomics (LFQ) employing normalised spectral abundance factors (NSAF) to determine the relative abundance of each protein to determine a ratio between SB treatment/DMSO vehicle [50, 51].

We identified 176 differentially upregulated and 137 differentially downregulated proteins (p < 0.05) in the SB treated cells compared to the DMSO vehicle control (Supp. Table 2). To determine cellular pathway changes upon SB treatment, Ingenuity Pathway Analysis (IPA) was used to predict activation and/or inhibition of canonical pathways and processes using experimental values of proteins from LFQ. IPA predicted the activation of protein kinase A signaling (Z-score = 0.832, p-value = 3.81−10^-4^) and AMPK signalling (Z-score = 0.816, p-value = 1.46 x 10^-3^, Figure. 6A).

**Figure 6.**
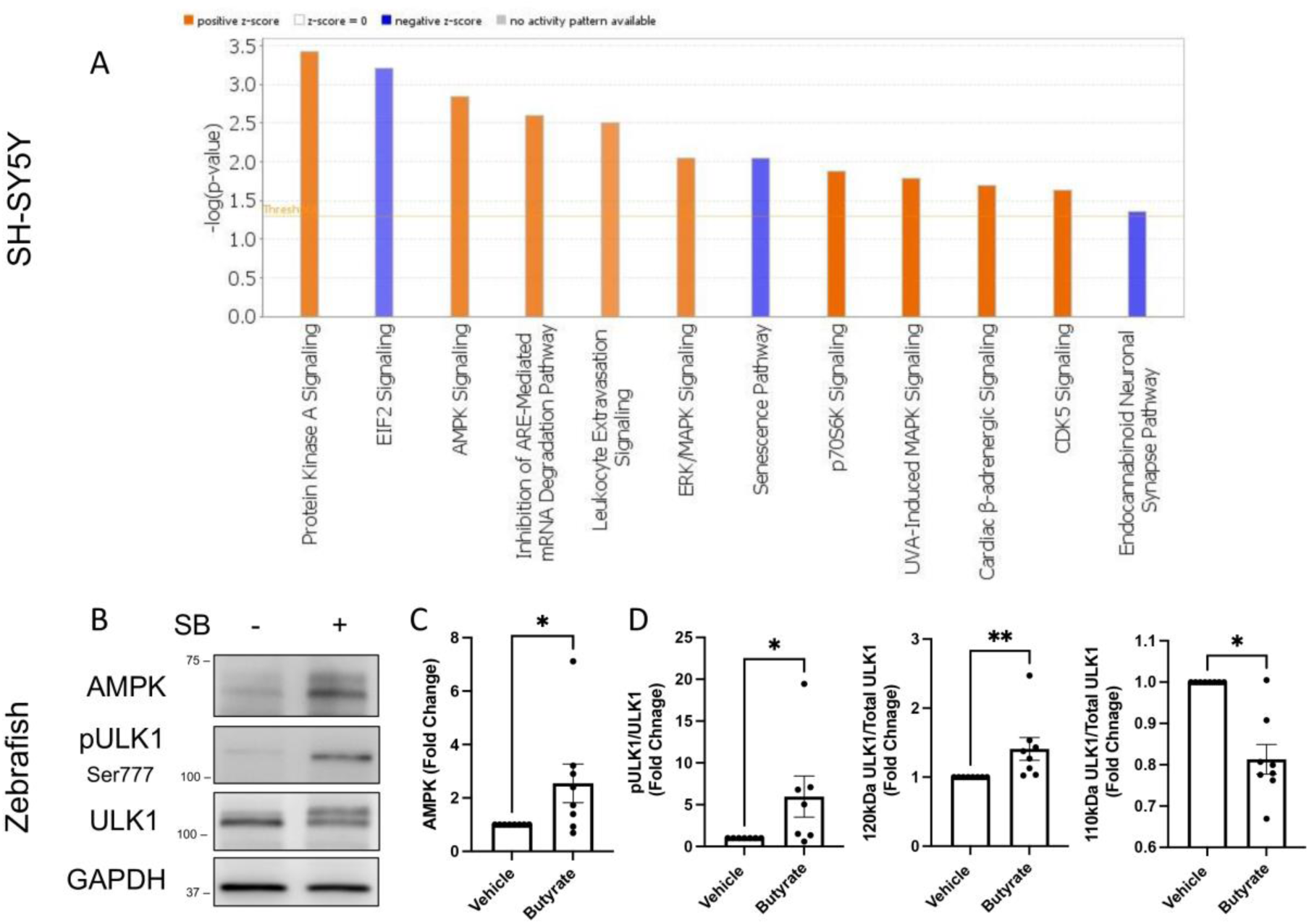
Proteomic analysis reveals upregulation of the Protein Kinase A and AMPK signaling pathway with SB treatment. (A) Ingenuity pathway analysis (IPA) predicted activation of the Protein Kinase A and AMPK signalling pathway upon SB treatment on ataxin-3 84Q expressing SH-SY5Y cells. Blue indicates IPA predicted an inhibition versus orange refers to an activation. (B) Immunoblots of ataxin-3 84Q zebrafish protein samples treated with SB or a vehicle control. Blots were probed with AMPK, phosphorylated ULK1 (Ser777) and ULK1. (C) Quantification revealed increased AMPK levels with butyrate treatment compared to vehicle treated (p = 0.0209, n=8). (D) Densitometric analysis revealed increased pULK1/ULK1 ratio in the SB treatment compared to the vehicle treated SCA3 zebrafish (p = 0.0469, n=7). Quantification of ULK1 revealed increased ULK1 protein at 120 kDa relative to total ULK1 and decreased 110kDa relative to total ULK1 in SB treated SCA3 zebrafish compared to the vehicle control (p = 0.0078 and p = 0.0156 respectively; n=8). Data represents mean ± SEM. Statistical analysis performed was a paired non-parametric t-test (Wilcoxon test).

To confirm the upregulation of the AMPK pathway, AMPK protein levels were analysed. SCA3 zebrafish treated with butyrate displayed a 2.5-fold increase in AMPK levels compared to the vehicle control (Figure. 6B, C; Uncropped images found in Supp. Figure 11). Similarly, levels of ULK1 were examined as it is a protein known to initiate autophagosome formation. Specifically, phosphorylation of ULK1 at position Ser777 by AMPK has been previously described to induce autophagy [52]. Western blotting detected phosphorylated ULK1 at Ser777 (pULK1) in the SB treated Ataxin-3 84Q zebrafish and densitometric analysis revealed approximately a 5x-fold increase compared to the vehicle treated control (Figure. 6B, D). Upon immunoblotting for ULK1, we noted two bands at 110 kDa and 120 kDa, whereby the 120 kDa band of ULK1 that was more marked in the SB treated Ataxin-3 84Q larvae. We have previously demonstrated that the pULK1 band overlies the 120kDa ULK1 band, suggesting that the 120kDa ULK1 band is likely pULK1 [49]. Quantification of these two ULK1 bands (relative to total ULK1) revealed greater amounts of the 120 kDa (1.3x), and decreased levels of the 110 kDa band (approximately 0.8x), in the presence of SB compared to the vehicle treatment, suggestive of autophagy induction (Figure. 6D).

To test whether the induction of autophagy by SB was dependent on AMPK activity we performed a co-treatment study using SB and a reported AMPK inhibitor GSK690693. We treated N2A cells expressing ataxin-3 84Q with bafilomycin, SB and GSK690693 and compared to those treated with bafilomycin and SB alone. Western blot analysis revealed an increase in the amount of LC3II/I present in the triple-treated ataxin-3 28Q cells (p=0.0156) and no difference in the triple-treated ataxin-3 84Q cells (p=0.0547) (see Supplementary Figure 9). These results suggest that although the proteomic data predicted that SB treatment is inducing autophagy through activation of AMPK, genetic silencing approaches would be required to demonstrate this conclusively.

## Discussion

### Sodium butyrate increases histone acetylation and decreased movement deficits in SCA3 zebrafish

We recently reported that our SCA3 zebrafish expressing EGFP-ataxin-3 84Q have higher levels of acetylated of histones 3 and 4 compared to non-transgenic control zebrafish at 6 days post fertilization [26]. This finding is consistent with ataxin-3 protein having a role transcriptional regulation [18, 21, 53] and previous reports of expression of polyQ expanded ataxin-3 resulting in histone hypoacetylation [22, 24–26]. Here we found that treatment with sodium butyrate prevented this effect on acetylation, increasing acetylation of both histones 3 and 4. This increase in the presence of acetylated histone 3 and 4 is likely caused by histone deacetylase (HDAC) inhibition effects induced by SB treatment. Along with the increase in histone acetylation, we also found that SB treatment improved the swimming of SCA3 zebrafish when compared to control zebrafish.

These findings contribute to existing evidence that suggests amelioration of histone hypoacetylation through HDAC inhibition can improve motor phenotypes in pre-clinical models of polyQ diseases [21, 23–27, 29, 35–38, 42, 54]. Other studies relating to the treatment of mouse models of SCA3 with SB are in alignment with our findings [22, 53]. The transgenic SCA3 mice developed by Chou and colleagues show a motor phenotype, neuropathology and hypoacetylation of histone 3 and histone 4. Chronic treatment of SCA3 mice with SB restored movement performance, to similar levels to wild type controls, with notable improvements in latency to fall off the rotarod, locomotor activity, foot dragging and dendritic arborisation [25].

### Sodium butyrate increases activity of the autophagy protein quality control pathway

In this study, we firstly identified that neuronal-like (neuroblastoma) SH-SY5Y cells expressing ataxin-3 containing a polyQ expansion (84 polyQ repeats) carried ataxin-3 protein aggregates. Interestingly, our study did not detect predominately nuclear ataxin-3 protein aggregates, which differs to some previous reports of polyQ expanded ataxin-3 resulting in nuclear protein aggregates [55].

In our study we found that 3-day treatment of SCA3 SH-SY5Y cells and 5-day treatment of transgenic SCA3 zebrafish with SB both resulted in removal of the protein aggregates (detected by confocal imaging), decreased presence of uncharacterised HMW ataxin-3 species and decreased the number of insoluble ataxin-3 particles detected by flow cytometric (FloIT) analysis. In contrast with these findings, we did find that shorter periods of SB treatment (24 hours in SH-SY5Y cells and transgenic SCA3 zebrafish) resulted in either an increase in the number of detergent insoluble ataxin-3 particles, or no significant effect. We hypothesise that the initial increase in insoluble particles observed may be a consequence of HDAC inhibition, causing increased transcription and translation of mutant ataxin-3 protein. In contrast, more prolonged treatment with SB (3-5 days) produced a robust decrease in ataxin-3 proteinopathy, demonstrating removal and degradation of ataxin-3 protein species. In view of this, prolonged administration of SB may be required to increase autophagic activity and yield optimal therapeutic benefit.

We also identified that SB treatment was inducing autophagy, indicated by an effect of co-treatment with autophagy inhibitors, 3MA, bafilomycin A1 or chloroquine, on LC3II abundance, both *in vitro* and *in vivo*. We likewise found that the decrease in HMW ataxin-3 bands produced by SB treatment was autophagy-dependent. These findings align with a previous report by Qiao et al [56] that SB treatment induced autophagy in enteroendocrine cells expressing α-synuclein. Combination treatment of SB with trehalose, a disaccharide, within a rodent model of Parkinson’s disease revealed increased activation of autophagy alongside reduction of pre-fibrillar form of phosphorylated α-synuclein [57]. Nevertheless, our findings are the first evidence of SB removing protein aggregates in a neuronal-like cell line and *in vivo,* presumably through the autophagy pathway.

Interestingly, within this study we have found positive effects of SB treatment, on zebrafish swimming, despite increased abundance of monomeric ataxin-3 protein resulting from the treatment. This finding is in line with other studies using the same zebrafish model, wherein increased swimming has been present despite increased human ataxin-3 protein, likely due to other protective effects [26]. Multiple protective mechanisms have been proposed to result from treatment with compounds with HDAC inhibitor activity. These include regulation of transcription, and in turn increased plasticity, neuronal regeneration, increased levels of neurotrophic factors such as GDNF and BDNF, restoration of balance to cerebellar circuitry and even protection against glutamate-induced excitotoxicity [22, 27, 57–59] . Our findings demonstrate that induction of autophagy is another therapeutic benefit of treatment with the HDAC inhibitor compound SB. Multiple other HDAC inhibitor compounds, such as sodium valproate and resveratrol, have similarly been demonstrated to induce autophagy [26, 60–64].

To understand the mechanism by which SB was inducing autophagy we performed proteomic analysis on lysates from SH-SY5Y cells expressing human ataxin-3 84Q treated with SB or vehicle control. Label-free proteomics analysis and Ingenuity Pathway Analysis predicted that SB treatment was activating protein kinase A and AMPK signalling. We found through validation by immunoblot analysis that indeed, SB treatment in the SCA3 zebrafish triggered increased levels of autophagy related protein AMPK and a downstream substrate phosphorylated ULK1 (Ser777). Whilst increased AMPK activity following SB exposure has been previously reported in colorectal and bladder cancer treatment studies [65–67], it has not been reported in neurodegenerative disease treatment studies. To test whether the induction of autophagy by SB was dependent on AMPK activity we performed a co-treatment study using SB and a reported AMPK inhibitor GSK690693. We did not detect a decrease in the autophagy induction produced by SB (indicated by increased LC3II in bafilomycin treated cells), but instead found no difference or an increase in LC3II/I in treated cells, likely due to an effect of this compound on the activity of a range of other kinases.

Protein kinase A and AMPK signalling commonly affect similar proteins such as FOXO1. Increased activation of FOXO1 is known to induce autophagy following treatment with HDAC inhibitors by increasing transcription of autophagy related genes (*Atg4b, Atg12, Pik3c3, Becn1*, and *Map1lc3b*) [68]. It has been reported that AMPK activates FOXO1 via phosphorylation of site Thr694 [69]. Interestingly, the FOXO family of transcription factors have been reported to have pro-longevity effects [70, 71] and are reported to be down regulated with aging [72], suggesting a potential benefit of FOXO1 related therapeutics in age-linked neurodegenerative diseases [73]. Most recently, it has been demonstrated that treatment with ketone body, D-β-hydroxybutyrate, can activate the autophagy pathway through the AMPK pathway and activate transcription factors FOXO1 and FOXO3a in healthy rat cortical neurons [74]. Previously, overexpression of DAF-16, the equivalent gene of FOXO in *C. elegans*, prevented motor impairment in a *C. elegans* model of SCA3 [75]. From these studies, alongside our results, the relationship between AMPK-mediated autophagy induction and downstream activation of FOXO transcription factors warrants further investigation as a potential target for SCA3 treatment.

There are several studies that describe impaired autophagy dynamics within SCA3 patient brains and multiple *in vitro* and *in vivo* models of SCA3 [45–48, 76, 77]. These studies have demonstrated decreased expression of autophagy related genes in SCA3 patients, the sequestration of proteins involved in autophagy dynamics within aggregates found in SCA3 patient brain samples, and modulated autophagosome production [45–48, 76, 77]. Our team recently found impaired autophagy dynamics within SH-SY5Y cellular and transgenic zebrafish models of SCA3 [49]. Collectively, these factors suggest that deficits in autophagy function may contribute to ataxin-3 proteinopathy. Several studies have also demonstrated that induction of the autophagy pathway, via genetic modification [11, 45, 76, 78] or exposure with small compounds [44, 79–85], can have beneficial effects as a treatment of SCA3 models. For example, trehalose has been tested across *in vitro* and *in vivo* models of SCA3 and has shown to decrease ataxin-3 positive aggregates [77], increase cerebellum layer thickness [86], and improve motor function [49, 86]. Moreover, these pre-clinical studies led to a clinical trial for safety and efficacy in SCA3 patients with minor adverse events and stabilisation of disease progression [87].

The findings within this study have relevance to a broad range of neurodegenerative diseases as there is growing evidence to suggest that changes to the gut microbiome, and therefore changes to levels of metabolites released by the gut, such as butyric acid, may impact the function of the nervous system [28, 56, 88, 89]. This is because, whilst these metabolites are naturally produced in the gut, they may elicit effects in the CNS through either the gut-brain axis or through crossing the blood brain barrier from the circulation [28, 56]. Our team has recently investigated whether the gut microbiome is altered in a SCA3 mouse model and found that indeed SCA3 mice have an altered gut microbiota structure and composition, including a decreased abundance of some populations of butyrate-producing bacteria [90]. Therefore, butyrate-related treatments, or exploiting the gut microbiome to increase butyrate production, may have therapeutic potential for SCA3. Whilst not explored within a SCA3 model, a recent study found that placing healthy mice on a ketogenic (low carbohydrate, high fat) diet resulted in increased levels of D-β-hydroxybutyrate in the blood and activation of the autophagy pathway [74]. Further, combined treatment of taurursodiol, a soluble form of bile acid, and phenylbutyrate, a derivative of butyric acid, has been found to improve survival in clinical cohorts of ALS patients [41] and received FDA approval for the treatment of ALS.

In contrast with our findings, previous studies investigating the therapeutic potential of SB in neurodegenerative models, including a mouse model of Huntington’s disease [42] and an *in vitro* model of Parkinson’s disease [56], found no effect of SB treatment on the number of protein aggregates present within those models. However, one report of butyrate treatment of a SOD1G93A mouse model of amyotrophic lateral sclerosis did report a reduction in the number of SOD1 positive aggregates in the colon, compared to vehicle treated animals [91]. Despite this, the authors did not report whether SOD1 protein aggregates were reduced within neurons, or whether the autophagy pathway was induced by SB treatment. One explanation for why this finding has not been reported previously is that the optimal dose (and dosing frequency) required for SB to induce autophagy may fall within a relatively narrow range, or it may vary depending on specific protein quality control impairments induced by the individual gene mutations. Doses reported within this study varied from previous doses reported to ameliorate neuropathology in a mouse model of SBMA [23], or in a transgenic mouse model of Huntington’s disease [42]. Further studies are required to confirm the optimal dosage at which SB can induce degradation of ataxin-3 protein aggregates in neurons, as the dosage required may vary between different experimental animal models of SCA3.

### Concluding Remarks

Sodium butyrate treatment was found to affect the aggregation status of polyQ expanded ataxin-3 *in vitro*, decreasing presence of high molecular weight (HMW) ataxin-3 bands and enhancing degradation of ataxin-3 positive aggregates. This study also provides the first experimental evidence of sodium butyrate treatment inducing increased autophagic activity via the PKA/AMPK pathways within a model of a neurodegenerative disease. Induction of the autophagy protein quality control pathway may be a therapeutic option for neurodegenerative diseases due to the potential to increase the clearance and degradation of toxic protein species such as mutant ataxin-3 protein. It must also be considered that there may also be limitations to the benefit of autophagy inducing candidates for patient treatment. A previous trial of treatment with lithium carbonate, known to induce autophagy, did not report positive outcome measures [92].

We propose that our finding that treatment with SB can induce autophagy, both *in vitro* and *in vivo*, resulting in protective effects such as preventing the development of impaired movement are important for research into treatments for SCA3. These results indicate that sodium butyrate and butyric acid both warrant further investigation as autophagy inducing agents with therapeutic potential for a wide range of neurodegenerative and protein aggregation diseases.

## Methods

### Transgenic SCA3 zebrafish

All animal experiments were performed in accordance with the Animal Ethics Committee of Macquarie University, N.S.W., Australia (ARA: 2016/004 and 2017/019). Adult zebrafish (*Danio rerio*) were housed in a standard recirculating aquarium system at 28.5 °C on a 14:10 light:dark cycle with twice daily feeding of artemia and standard pellet [93]. These experiments used our previously described transgenic zebrafish model of SCA3 [44]. Briefly, the driver lines express a HuC (*elavl3*) neuronal promoter driving Gal4 VP16 [Tg(elavl3:GAL4-VP16-mCherry)mq15], and the responder lines express EGFP-ATXN-3 (containing 23 or 84 CAG repeats) and dsRED under the control of the UAS promoter that is activated only in tissues that are expressing Gal4 VP16 [Tg(UAS:Hsa.ATXN3_23xCAG-EGFP,DsRed)mq16 and Tg(UAS:Hsa.ATXN3_84xCAG-EGFP,DsRed)mq17]. For the studies described here driver and responder lines were mated to generate F1 HuC-EGFP-Ataxin-3 23Q or 84Q lines, which were in-crossed to generate F2 embryos for use in this study.

### SCA3 cell culture model

SH-SY5Y cells were grown in Dulbecco’s Modified Eagle’s Medium (DMEM)/Nutrient Mixture F12 Ham supplemented with 10 % fetal bovine serum (FBS) and maintained at 37 °C and 5 % CO_2_. pcDNA3-myc-Ataxin3Q28 and pcDNA3-myc-Ataxin3Q84 were a gift from Henry Paulson (Addgene plasmids # 22124 and #22125, unmodified in house); [94]. These vectors, containing full length ataxin-3 (28Q or 84Q) and a neomycin resistant gene, were used for stable selection of transfected cells.500 µg/mL of neomycin was used to select cells stably expressing ataxin-3. Cells were treated with 250 µg/mL of neomycin (Sigma) to maintain expression of the selective expression of the ataxin-3 protein or vector control.

### Drug treatments in SCA3 cell cultures and transgenic SCA3 zebrafish

SH-SY5Y cells stably expressing human ataxin-3 were seeded into a 24-well plate at a density of 40,000 cells/cm^2^ and incubated at 37 °C supplemented with 5 % CO_2_. 24 hours after seeding, cells were treated with SB (3 mM, Cayman Chemicals [Cat #13121]) or vehicle control (autoclaved water) diluted in growth media for a total of 72 hours, with SB-containing media replenished every 24 hours to ensure drug efficacy. For the co-treatment study with an autophagy inhibitor, cells were pre-treated with 3MA (5 mM, Cayman Chemicals) for 1 hour before SB was added and left on for the duration of the SB treatment or bafilomycin A1 (100 nM dissolved in DMSO, Roche) was added for 4 hours of prior to protein extraction.

For the zebrafish drug treatment studies, zebrafish embryos (1 day post fertilization; dpf) were screened for fluorescence (EGFP and dsRED) indicating that they were positive for the EGFP-ataxin-3 transgenes. In the drug treatment groups, positive embryos were treated with a single administration for five days with SB (250 µM, 500 μM and 1 mM, solubilized in milliQ water). Appropriate volumes of SB were diluted in zebrafish E3 medium, with the control group containing only E3 media [95]. Co-treatment with SB and the autophagy inhibitor chloroquine (1.5 mM, Sigma-Aldrich [Cat# C6628], solubilized in milliQ water) was performed in the same manner but with chloroquine stock solution added to the E3 media containing SB so that the solution also contained 1.5 mM chloroquine. Vehicle treated groups were left in E3 medium. Approximately 25-30 embryos were treated per group per experiment.

### Zebrafish motor behavior testing

Motor function assays were performed as described in Watchon et al [44]. All behavioural tracking was performed using a ZebraLab Tracking System (ZebraBox; Viewpoint). Tracking of 6 dpf larvae were conducted by randomly assorting them in 24 multi-well plates within the ZebraBox housed with an enhanced light source, under conditions of 6 minutes light, 4 minutes dark and 4 minutes light. The total distance travelled by each larva within the dark phase was calculated.

### Western blotting

SH-SY5Y cells stably expressing human ataxin-3 (28Q and 84Q), were washed with ice cold PBS and incubated in 100 µL of RIPA buffer containing protease inhibitors (Complete ULTRA tablets, Roche). Cells were gently agitated on orbital shaker for five minutes before being spun down at 18 000g for 15 minutes. Protein lysates were prepared from zebrafish larvae (6 dpf) in RIPA buffer containing protease inhibitors (Roche), using a manual dounce homogenizer. Homogenates were centrifuged for 20 minutes at 15 000g. Supernatants were collected and measured for protein concentration using the Pierce™ BCA Protein Assay Kit (Thermo Fisher Scientific).

Equal amounts of protein were separated using SDS-PAGE and transferred to PVDF membrane for immunoblot probing. Antibodies used included anti-rabbit ataxin-3 (kind gift from H. Paulson) and anti-mouse GAPDH (Proteintech). To test the effect of drug treatments, immunoblots were probed for anti-rabbit acetylated histone 3 at lysine 9 and histone 4 at lysine 5 (ac-H3K9 and ac-H4K5 respectively, Cell Signaling), anti-rabbit LC3B (Abcam), anti-rabbit AMPK (Cell Signaling), anti-rabbit phosphorylated-ULK1 (Ser777; Thermo Fisher), anti-rabbit ULK1 (Cell Signaling), and anti-mouse beta-actin (Sigma). The immunoblots were probed with appropriate secondary antibodies (Promega and Li-Cor) and visualised by chemiluminescence (Supersignal detection kit, Pierce) using a BioRad GelDoc System or fluorescent detection using the LiCor Odyssey CLX. The intensity of bands within the immunoblot were quantified by Image Studio Lite and the target protein expression level was determined by normalizing against the loading control protein.

### Preparation of Cultured SH-SY5Y cells for flow cytometry

For flow cytometry experiments, SH-SY5Y cells were transiently transfected with human ATXN3 plasmids containing a fluorescent tag. Human ATXN3 cDNA was subcloned into a pCS2+ vector to generate ATXN3 constructs containing a short polyQ (28Q) and an expanded polyQ (84Q) fluorescently tagged with EGFP.

SH-SY5Y cells were cultured in DMEM/Ham’s Nutrient Mixture F12 and supplemented with 10 % FBS and 5 % CO_2_. Cells were seeded into 6-well plates and incubated at 37 °C. Once cells had reached 85 % confluency (24-48 hours), cells were transiently transfected with 1 µg of DNA/3.5 cm^2^ using Lipofectamine 2000 (ThermoFisher, Catalogue # 11668030). Cells were treated with vehicle or 3 mM SB for 24 hours.

To determine transfection efficiency, live cells were imaged within 6-well plates at 20x magnification using an EVOS FL monochrome microscope (Invitrogen, catalogue #AMF4300) running Invitrogen image acquisition software. Cells were harvested using 0.5 % trypsin/EDTA and pelleted (1500 rounds per minute, 5 minutes at room temperature [RT]). Cells were re-suspended in 500 µL of lysis buffer (PBS containing 1 % Triton-X 100 and complete protease inhibitors [Roche]). DAPI was added to lysis buffer (final concentration 5 µM) and incubated at RT for 5 minutes, protected from light.

### Dissociation of zebrafish for flow cytometry

Whole zebrafish larvae (2 or 6 dpf) were euthanised and larvae were digested and dissociated into a single cell solution, as previously described [43]. In brief, following euthanasia, whole larvae were dissected into smaller pieces using a scalpel and forceps, and enzymatically digested using 0.5 % Trypsin-EDTA. Dissociated cells were then lysed using PBS containing 0.5 % Triton-X 100 and a marker of nuclei, either DAPI (final concentration 5 μM) or 1 x Red Dot (Gene Target Solutions, catalogue # 40060).

### Flow cytometric analysis of insoluble ataxin-3

Flow cytometry was performed using a Becton Dickinson Biosciences LSR Fortessa analytical flow cytometer and calibrated using CST beads (Becton Dickinson). The fluorescence of transfected cells was compared to an un-transfected control sample. Lysed samples were analysed by plotting forward scatter (area) against relative DAPI (379-28-A) or EGFP (530-30-A/535) fluorescence. Forward scatter threshold was set at 200 to minimize exclusion of small insoluble particles [96]. A total of 20 000 events were acquired for cell culture experiments and 50 000 events for dissociated zebrafish experiments. All axes were set to log_10_. Nuclei were identified and gated based on intensity of UV fluorescence and relative size (forward scatter). The number of insoluble GFP particles, indicating insoluble ataxin-3 particles, was analyzed based on GFP fluorescent intensity and forward scatter [43]. Gating of Triton-X insoluble GFP^+^ positive particles was performed by comparing populations to an un-transfected control sample. For analysis comparing the effect of different drug treatments on SCA3 cells and zebrafish expressing ataxin-3-84Q, data was analyzed as the number of insoluble GFP^+^ particles divided by the number of detected nuclei [43]. Drug-induced effects were presented as the fold change differences when compared to vehicle treated controls [43]. All flow cytometric analysis and gating was performed using FlowJo (version 10.6.2, Becton Dickinson Biosciences).

### In-gel Trypsin Digestion and Liquid Chromatography Mass spectrometry (LC-MS)

SH-SY5Y cell lysates expressing ATXN3-84Q were separated in 4-12 % NuPAGE Bis-Tris polyacrylamide gels (Invitrogen) in duplicate gels. One gel was Coomassie stained and de-stained in 25 % methanol while the second gel was transferred to nitrocellulose membrane and western blot analysis for ATXN3 was performed. Fraction 1-7 in the Coomassie stained gel were identified by comparison with the molecular weight of the ataxin-3-84Q band from the western blot results. The bands in the gel were excised and cut into 1-2 mm pieces and de-stained in 50 mM ammonium bicarbonate/50 % methanol pH 8, followed by 50 mM ammonium bicarbonate/50 % acetonitrile pH 8. The de-stained gel pieces were dehydrated in 100 % acetonitrile and the solution was removed to allow gel pieces to air dry. The proteins were reduced and alkylated with 10 mM dithiothreitol (DTT) and 20 mM iodoacetamide (IAA) respectively and digested with trypsin : protein [1:50 (w/w)] overnight at 37 °C as described [97].

Following overnight digestion, the supernatant was transferred to a fresh tube and the tryptic peptides from the gel pieces were extracted twice with 50 % acetonitrile/2 % formic acid and combined with the supernatants. The tryptic peptides were vacuum centrifuged to remove acetonitrile and desalted on a pre-equilibrated C_18_ Omix tip. The eluted peptides were further dried under vacuum centrifugation. The peptide pellet was the resuspended in 12 µL of 0.1 % formic acid and 10 µL was loaded for LC-MS/MS analysis.

Tryptic peptides were separated using an UHPLC Dionex Ultimate 3000 RSLC nano (ThermoFisher, USA) equipped with a Thermo Acclaim™ PepMap™ 100 C_18_ column (75 µm diameter, 3 µm particle size, 150 mm length) employing a 60 min gradient (2 %–26 % v/v acetonitrile, 0.1 % v/v formic acid for 40 mins followed by 50 % v/v acetonitrile, 0.1 % v/v formic acid for 10 mins and 80 % v/v acetonitrile, 0.1 % v/v formic acid for 8 mins) with a flow rate of 300 nl/min. The peptides were eluted and ionized into Q-Exactive Plus mass spectrometer (ThermoFisher, USA). The electrospray source was fitted with an emitter tip 10 μm (ThermoFisher, USA) and maintained at 1.6 kV electrospray voltage. FTMS analysis was carried out with a 70 000 resolution and an AGC target of 1−10^6^ ions in full MS (m/z range 400-2000); and MS/MS scans were carried out at 17 500 resolution with an AGC target of 2−10^4^ ions. Maximum injection times were set to 30 and 50 milliseconds respectively. A top-10 method was employed for MS/MS selection, ion selection threshold for triggering MS/MS fragmentation was set to 1−10^4^ counts, isolation width of 2.0 Da, and dynamic exclusion for 20 seconds was used to perform HCD fragmentation with normalized collision energy of 27.

Raw spectra files were processed using the Proteome Discoverer 2.4 software (Thermo) against the Swissprot database (organism *Homo sapiens,* version 25/10/2017 with 42252 sequences) incorporating the Sequest search algorithm. Peptide identifications were determined using a 20-ppm precursor ion tolerance and a 0.1 Da MS/MS fragment ion tolerance for FTMS and HCD fragmentation. Carbamidomethylation modification of cysteines was considered a static modification while oxidation of methionine, and acetyl modification on N-terminal residues were set as variable modifications allowing for maximum two missed cleavages. The data was processed through Percolator for estimation of false discovery rates. Protein identifications were validated employing a q-value of 0.01. For label-free quantitation, normalised spectral abundance factors (NSAF) were calculated for each sample according to Zybailov et al [51]. The relative fold-change of proteins from cells expressing ATXN-84Q were calculated by the ratio of SB treatment /DMSO vehicle control. The mass spectrometry proteomics data has been deposited to the ProteomeXchange Consortium via the PRIDE [98] partner repository with the dataset identifier PXD024626 and 10.6019/PXD024626. Username: and password: mZMbvai3.

### Data analysis

Data analysis was performed using GraphPad Prism software (Version 9). Group comparisons were made using one-way ANOVA tests, followed by a Tukey post-hoc to identify differences. Where data did not meet assumptions for one-way ANOVA, a non-parametric test, Kruskal Wallis, was used. Co-treatment studies were analysed using a two-way ANOVA (Factor 1: treatment with sodium butyrate, Factor 2: treatment with autophagy inhibitor, bafilomycin 1A/3MA or chloroquine). Flow cytometry experiments were analysed using student t-tests. Co-treatment data was analysed for main effects and interaction effects. Post-hoc multiple comparisons were used to determine statistical significance between treatment groups. Comparisons involving just two groups (e.g. vehicle and treated group) were analysed using a student t-test, or Wilcoxon test in cases of non-parametric data. All results presented are mean ± standard error mean (SEM) with statistical significance which is defined as *p≤0.05.

## Acknowledgements

The authors would like to thank Henry Paulson for the ataxin-3 antibody and EGFP-ATXN3 plasmids. We also thank staff members (past and present) of the zebrafish facility at Macquarie University for the care of the zebrafish described within this manuscript.

## Author contributions

MW, KJR, LL, EKD, GAN, AL, ASL designed the research; MW, KJR, LL, YA, KCY, SKP, FC, AL, ASL performed the research; MW, KJR, LL, YA, KCY, SKP, FC, AL, ASL analyzed data; MW, KJR, LL, YA, KCY, SKP, FC, EKD, AL, ASL interpreted and troubleshooted experimental results; MW, KJR, LL, YA, KCY, SKP, FC, EKD, AL, ASL wrote the paper; MW, KJR, LL, YA, KCY, SKP, FC, EKD, GAN, AL, ASL revised and approved the final submitted paper;

## Data availability

The data that support the findings of this study are available from the corresponding author upon reasonable request.

## Abbreviations

MJD: Machado-Joseph disease
SCA3: spinocerebellar ataxia type 3
HDAC: histone deacetylase
polyQ: polyglutamine
SB: sodium butyrate
dpf: days post fertilization
EGFP: enhanced green fluorescent protein
FSC: forward scatter signal
H3K9: acetylated histone 3 at lysine 9
H4K5: acetylated histone 4 at lysine 5

---

**Supplementary Figure 1.**
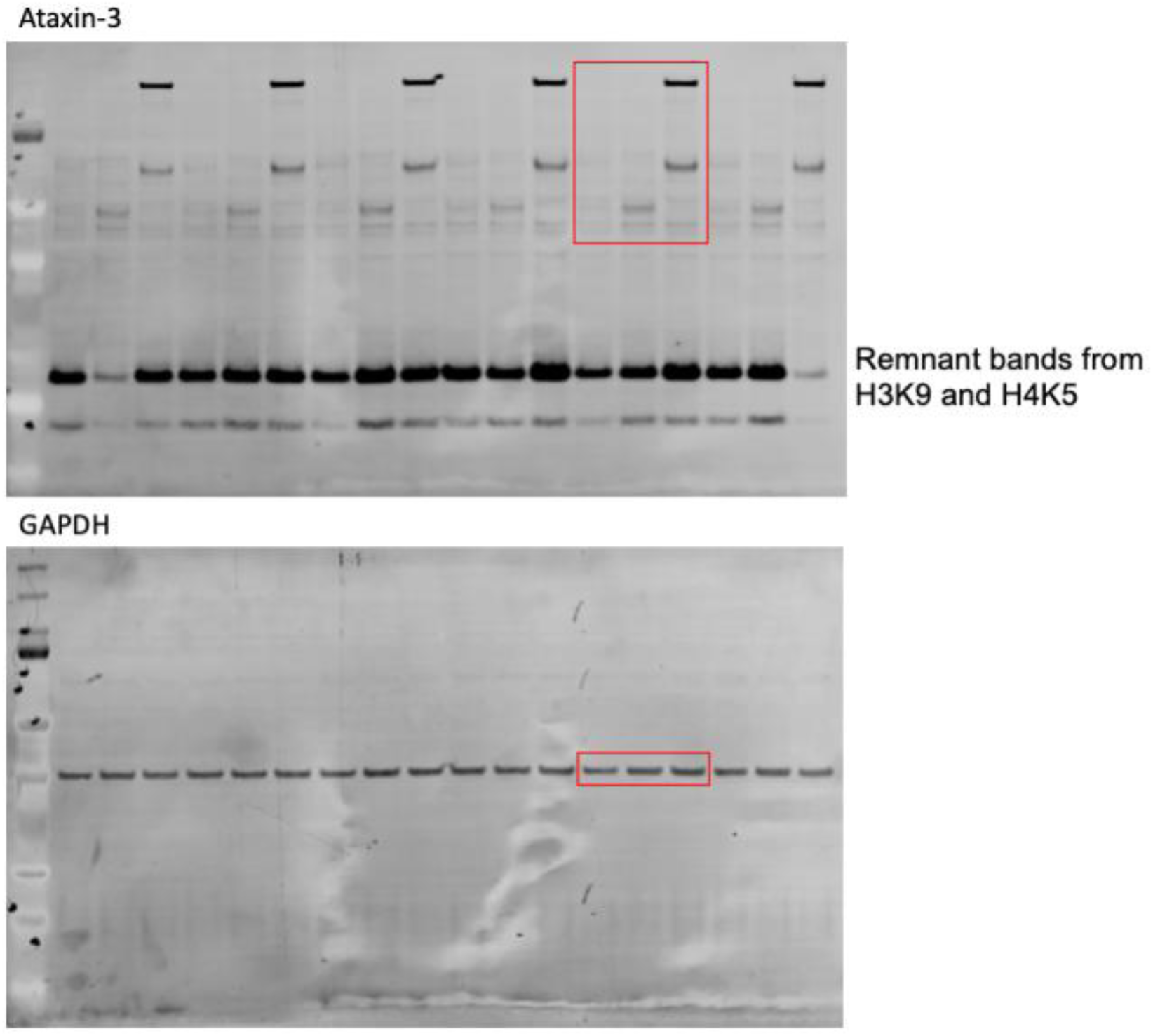
Uncropped western blot image of SH-SY5Y cells stably expressing human ataxin-3 (28Q and 84Q) with an empty vector control. Red box depicts representative image used in Figure 1A.

**Supplementary Figure 2.**
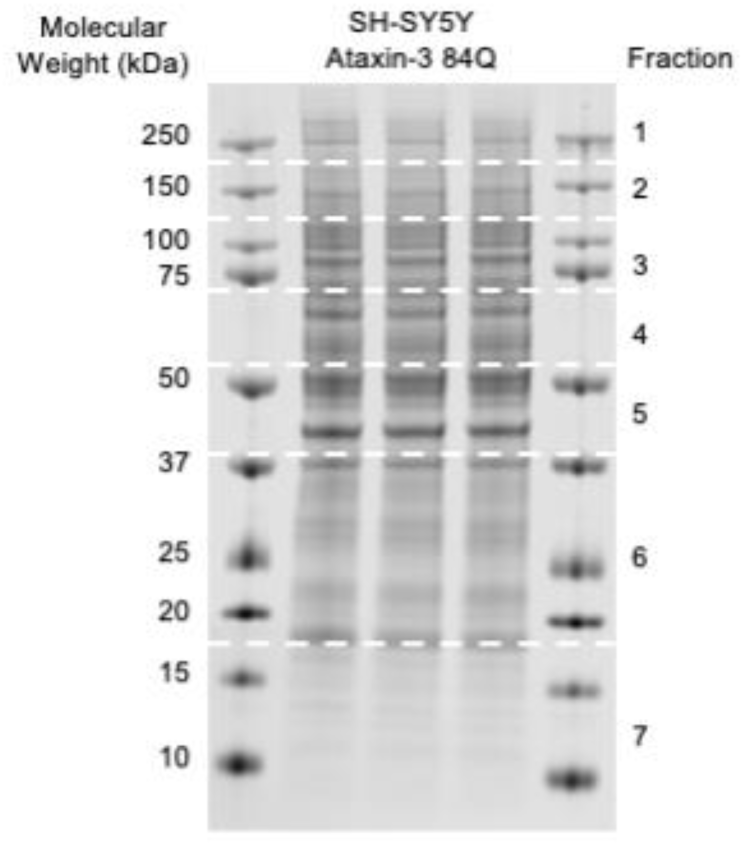
Validation of high molecular weight ataxin-3 expressed in ataxin-3-84Q SH-SY5Y cells using mass spectrometry. Representative SDS-PAGE detecting the total protein found in SH-SY5Y cells stably expressing ataxin-3 84Q. White dotted lines define the different fractions based on protein size for LC-MS/MS analysis.

**Supplementary Table 1. Identified ataxin-3 peptides in ataxin-3-84Q SH-SY5Y cells using mass spectrometry.**

Protein lysates of SH-SY5Y cells expressing ataxin-3-84Q were subjected to SDS-PAGE and different fractions were separated by size and subjected to using mass spectrometry aiming to identify the high molecular weight ataxin-3. Mass spectrometry identified ataxin-3 protein within Fraction 2, which contained proteins of between 120-200 kDa in size, where we predicted to find the high molecular weight ataxin-3.

**Supplementary Figure 3.**
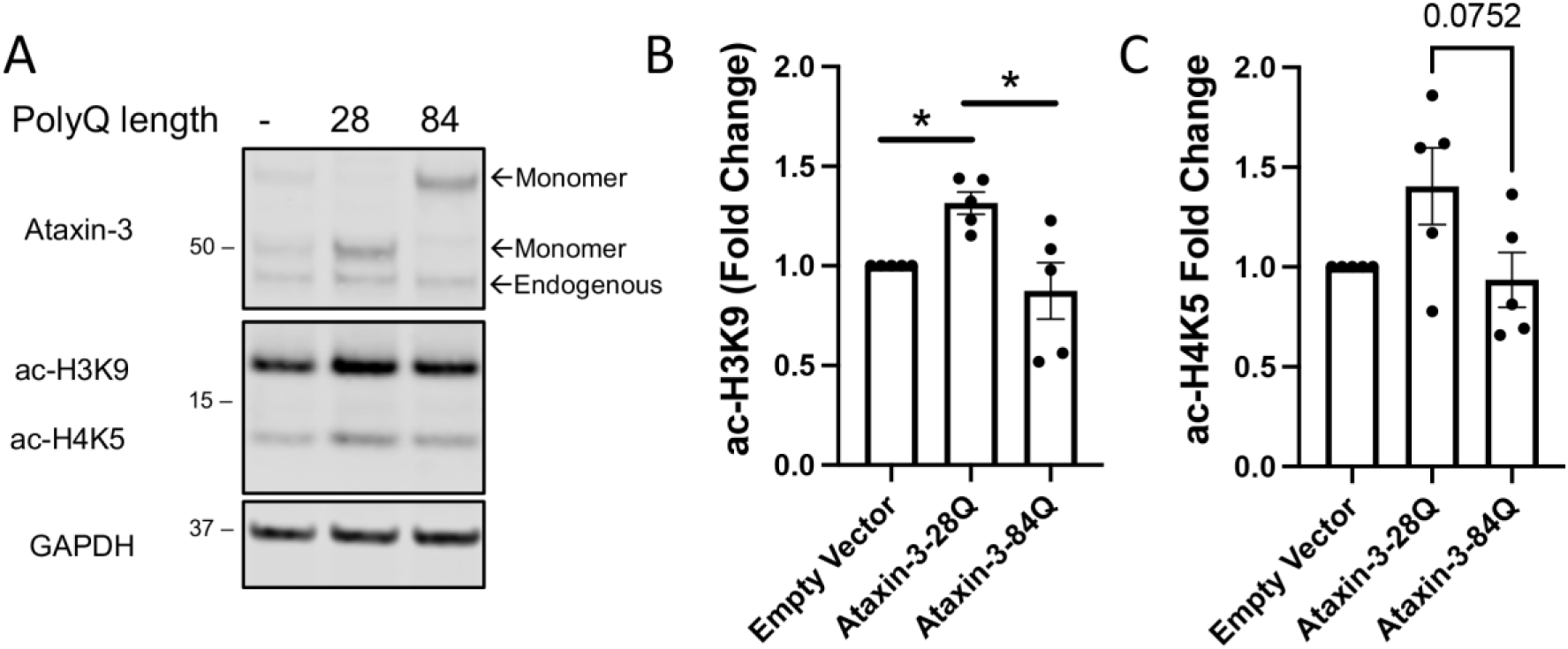
SH-SY5Y cells expressing ataxin-3 present with altered histone acetylation. (A) Western blot of stably expressing ataxin-3 28Q and 84Q SH-SY5Y cells comparing levels of ataxin-3, acetylated histone 3 at lysine 9 (ac-H3K9) and acetylated histone 4 at lysine 5 (ac-H4K5). (B) Densitometric analysis of ac-H3K9 revealed an increase in ac-H3K9 levels in ataxin-3 28Q cells compared to the vehicle control (p = 0.0431), whilst ataxin-3 84Q cells had decreased ac-H3K9 levels compared to the ataxin-3 28Q cells (p = 0.00186, n=5). (C) Quantifying the levels of ac-H4K5 were found to be similar between the different genotypes. Data represents mean ± SEM. Statistical analysis performed was a non-parametric one-way ANOVA, followed by Kruskal Wallis test.

**Supplementary Figure 4.**
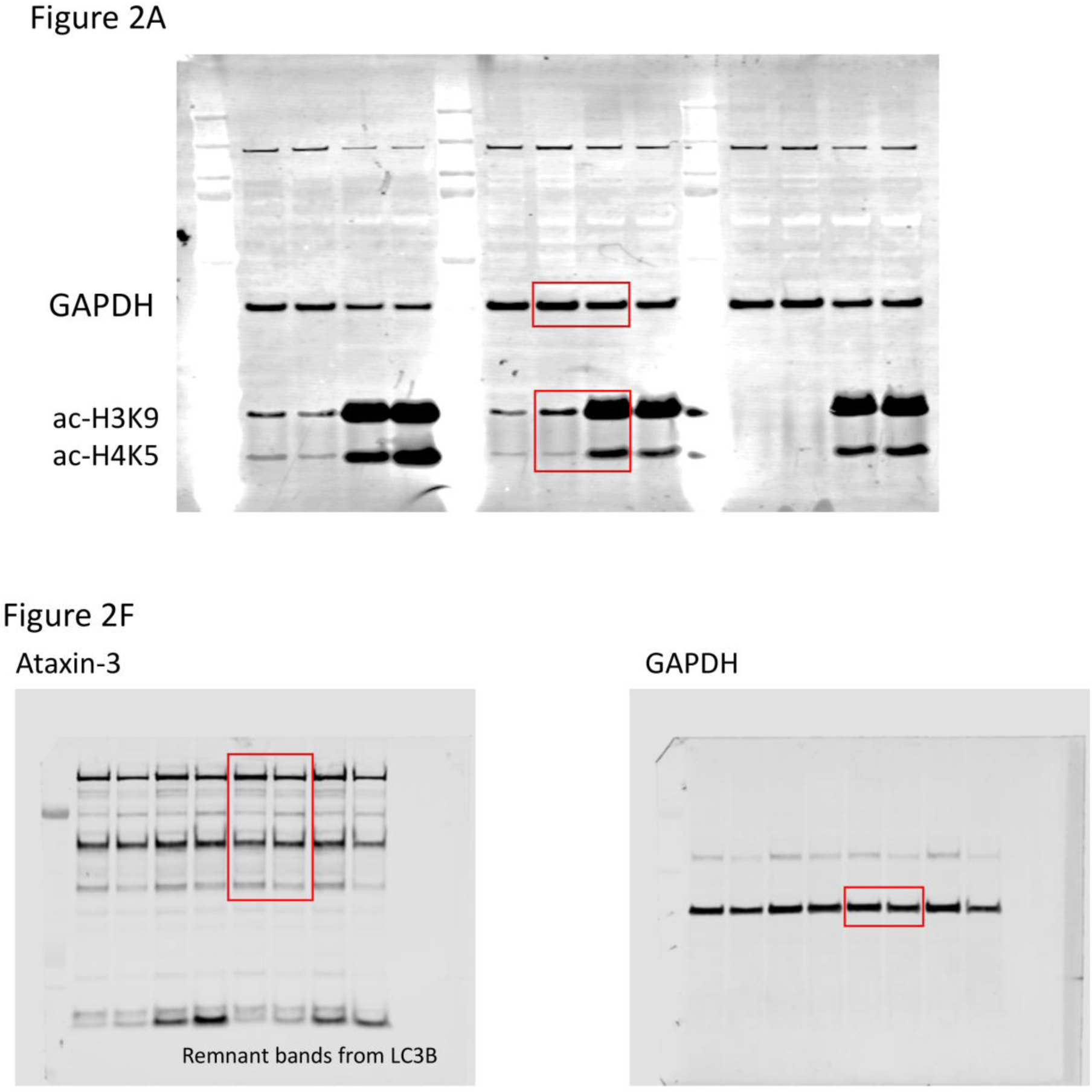
Uncropped western blot image of SCA3 cells treated with SB or vehicle treatment. Red box depicts representative images used in Figure 2A and Figure 2F.

**Supplementary Figure 5.**
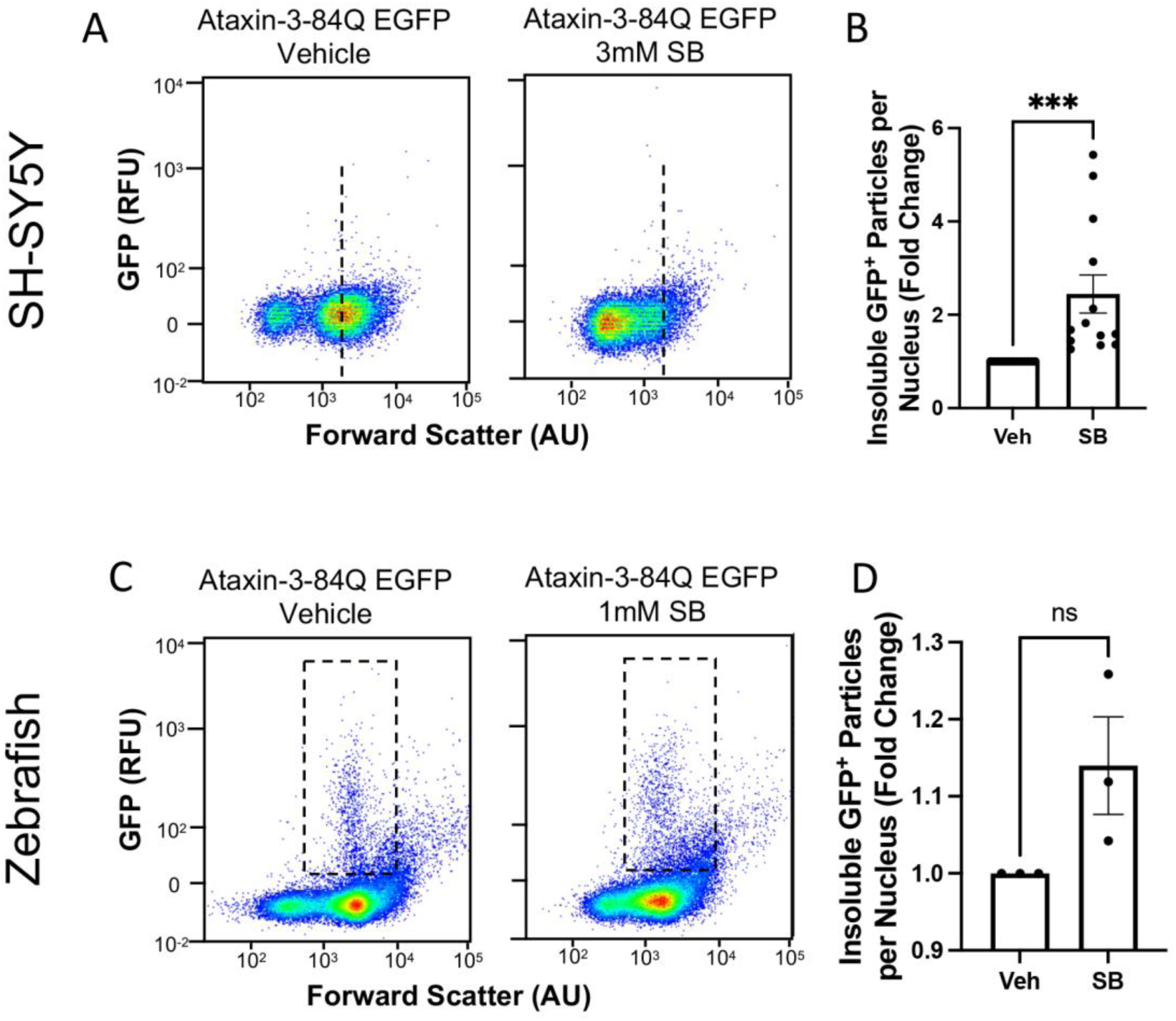
Short periods of SB treatment augments insoluble ataxin-3 aggregate load in SH-SY5Y cells and zebrafish expressing human ataxin-3 84Q. (A) Scatter plots representing the shift in size of EGFP^+^ Triton-X insoluble particles in SH-SY5Y cells following SB treatment. (B) SB treatment after 24 hours increased the number of GFP^+^ particles following flow cytometric analysis (p = 0.0002, n=7). (C) Representative scatter plots display Triton-X insoluble EGFP^+^ particles in two-day-old transgenic zebrafish expressing ataxin-3 with 84Q glutamine residues, with SB treatment producing a more disperse population of EGFP^+^ particles. (D) Quantification of detergent insoluble EGFP^+^ particles revealed an increase following SB treatment when compared to vehicle treatment, however this comparison did not reach statistical significance. Data represents mean ± SEM. Statistical analysis performed was an unpaired student t-test. Ns – non-significant.

**Supplementary Figure 6.**
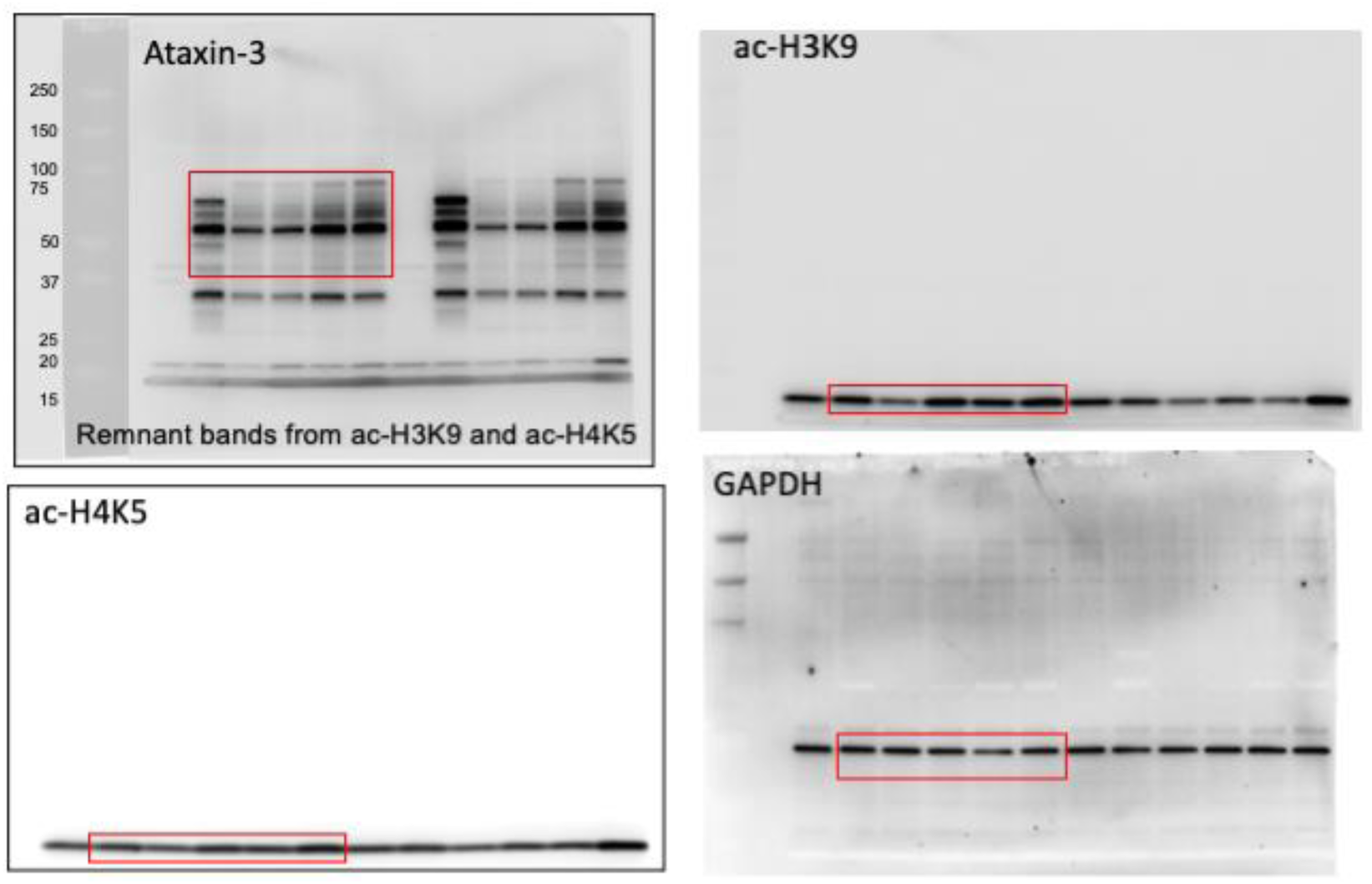
Uncropped western blot image of zebrafish expressing human ataxin-3 (23Q and 84Q) treated with SB. Red boxes depicts representative images used in Figure 3A.

**Supplementary Figure 7.**
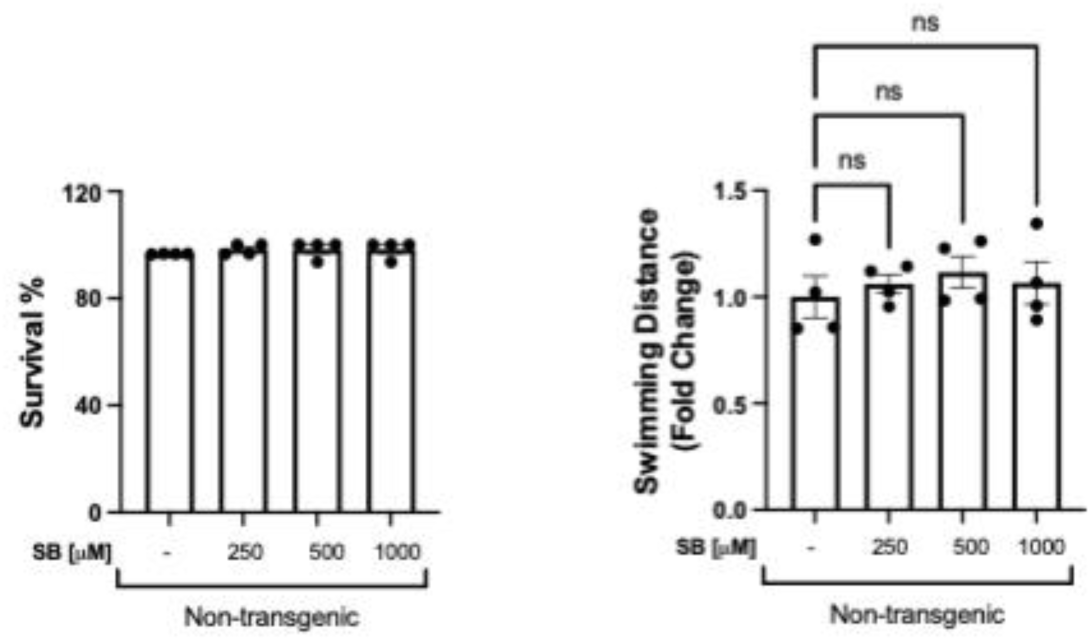
Survival and motor behaviour remains unchanged in non-transgenic zebrafish treated with sodium butyrate. (A) Non-transgenic zebrafish were treated with 250 μM – 1 mM SB between 1-6 days post fertilisation (dpf) and survival was calculated at 6 dpf. (B) Motor behaviour was analysed in non-transgenic zebrafish treated with SB at 6 dpf and found no differences between SB treatment compared to the control. Data represents Mean ± SEM. ns - non-significant. Data points reflect experimental replicates consisting of 25-30 embryos per replicate. Statistical analysis performed was a one-way ANOVA followed by a Tukey post-hoc test. Ns – non-significant.

**Supplementary Figure 8.**
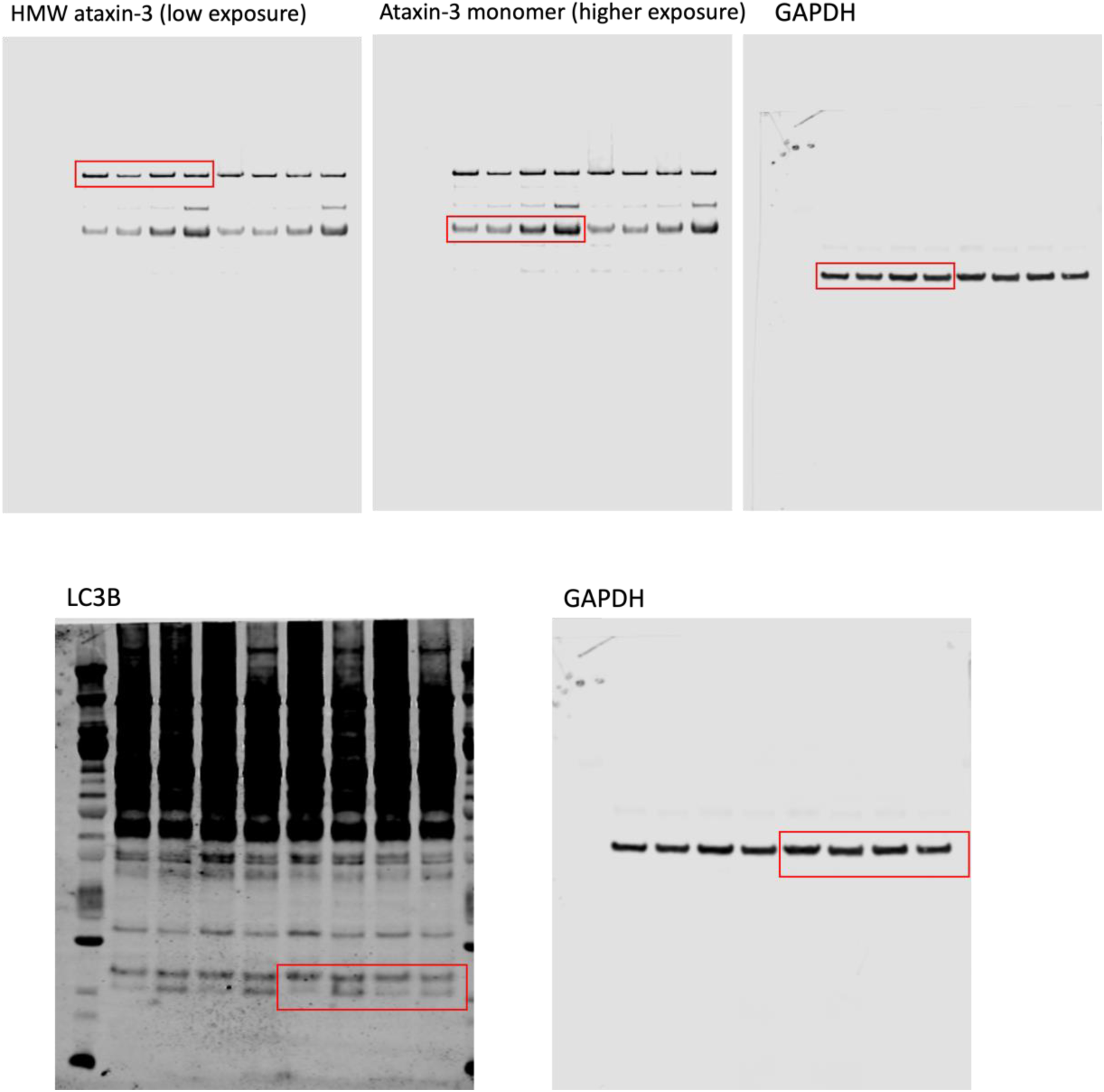
Uncropped western blot image of SH-SY5Y cells treated with either SB, 3MA or co-treatment. Red boxes depict representative images used in Figure 4A.

**Supplementary Figure 9.**
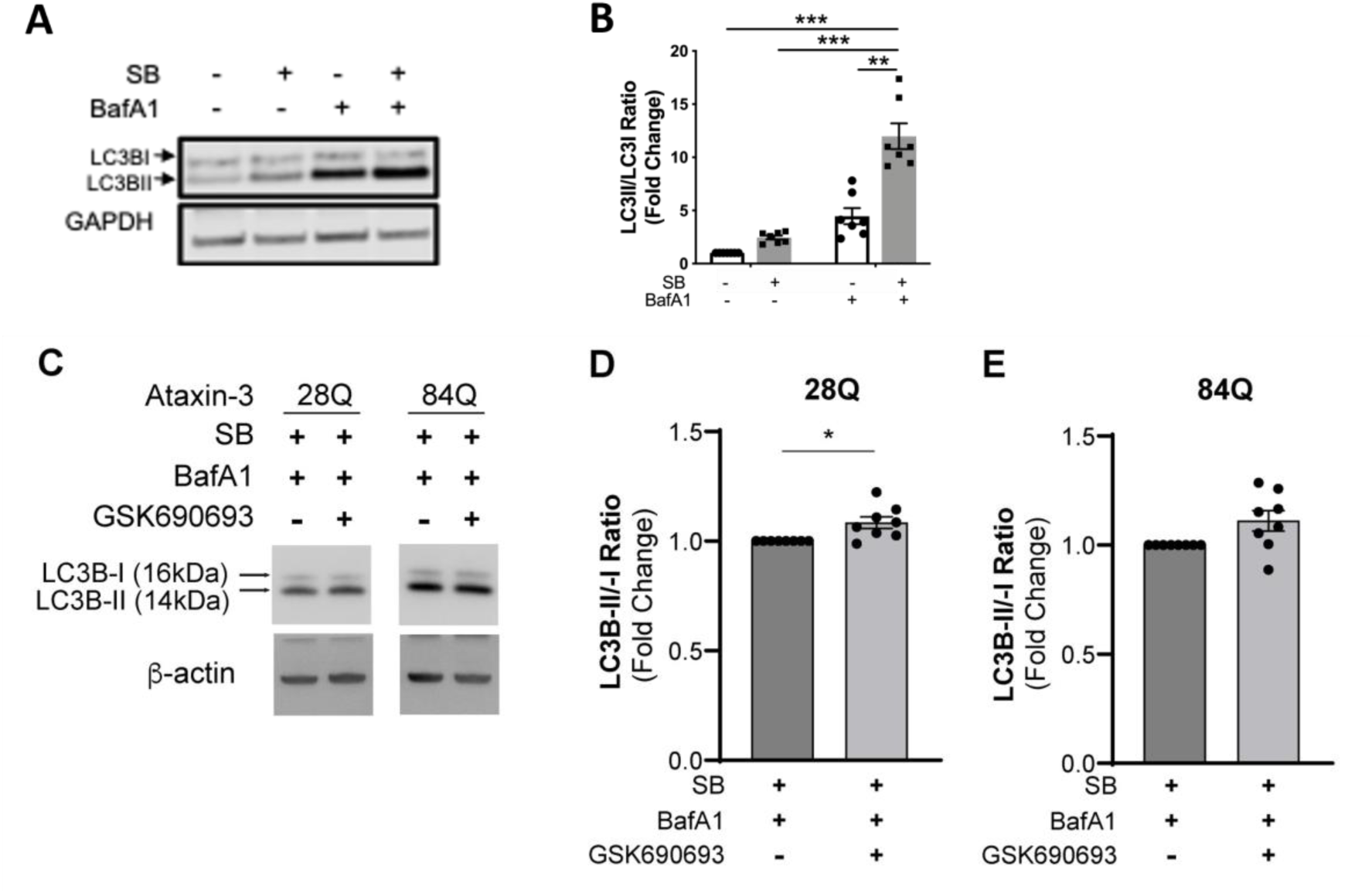
Sodium butyrate mediated autophagy induction is prevented by bafilomycin A1 co-treatment. (A) Western blot analysis was performed on lysates of SH-SY5Y cells stably expressing ataxin-3 84Q following treatment with 3 mM SB for 72 hours and/or 100 nM Bafilomycin A1 (Baf A1) for the final 4 hours, compared to a vehicle treatment. Blots were probed for LC3B. (B) Densitometry analysis of LC3B showed increased LC3II/LC3I ratio in SB and Baf A1 co-treated cells when compared to vehicle treatment (p < 0.0001), SB treatment (p < 0.0001) and Baf A1 treatment alone (p = 0.003, n=6-7). Statistical analysis performed was a two-way ANOVA followed by a Tukey post-hoc test. (C) Western blot analysis did not demonstrate a decrease in the amount of LC3II/I present in N2A cells expressing ataxin-3 28Q or ataxin-3 84Q treated with bafilomycin, SB and GSK690693, compared to those treated with bafilomycin and SB alone. (D) Densiometric analysis revealed that treatment with bafilomycin, SB and GSK690693, produced an increase in LC3II/1 compared to treatment with bafilomycin and SB alone, in ataxin-3 28Q cells (p=0.0156); and (E) no difference in ataxin-3 84Q cells (p=0.0547). Statistical analysis performed was Wilcoxon test due to non-parametric data. Data represents mean ± SEM.

**Supplementary Figure 10.**
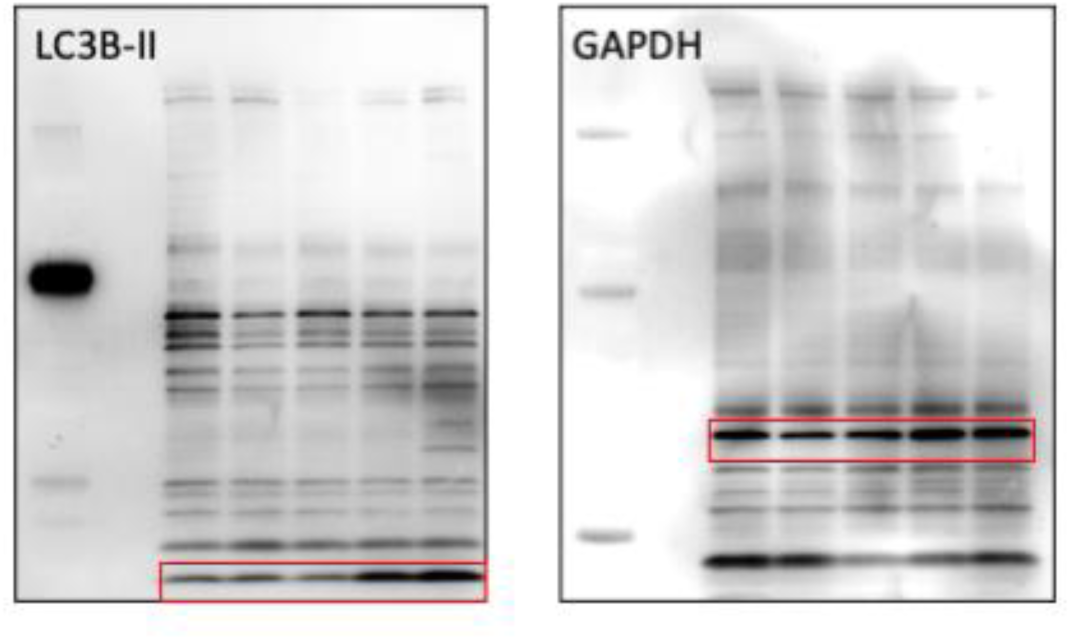
Uncropped western blot image of SCA3 zebrafish treated with either SB, chloroquine or co-treatment. Red boxes depict representative images used in Figure 5A.

**Supplementary Figure 11.**
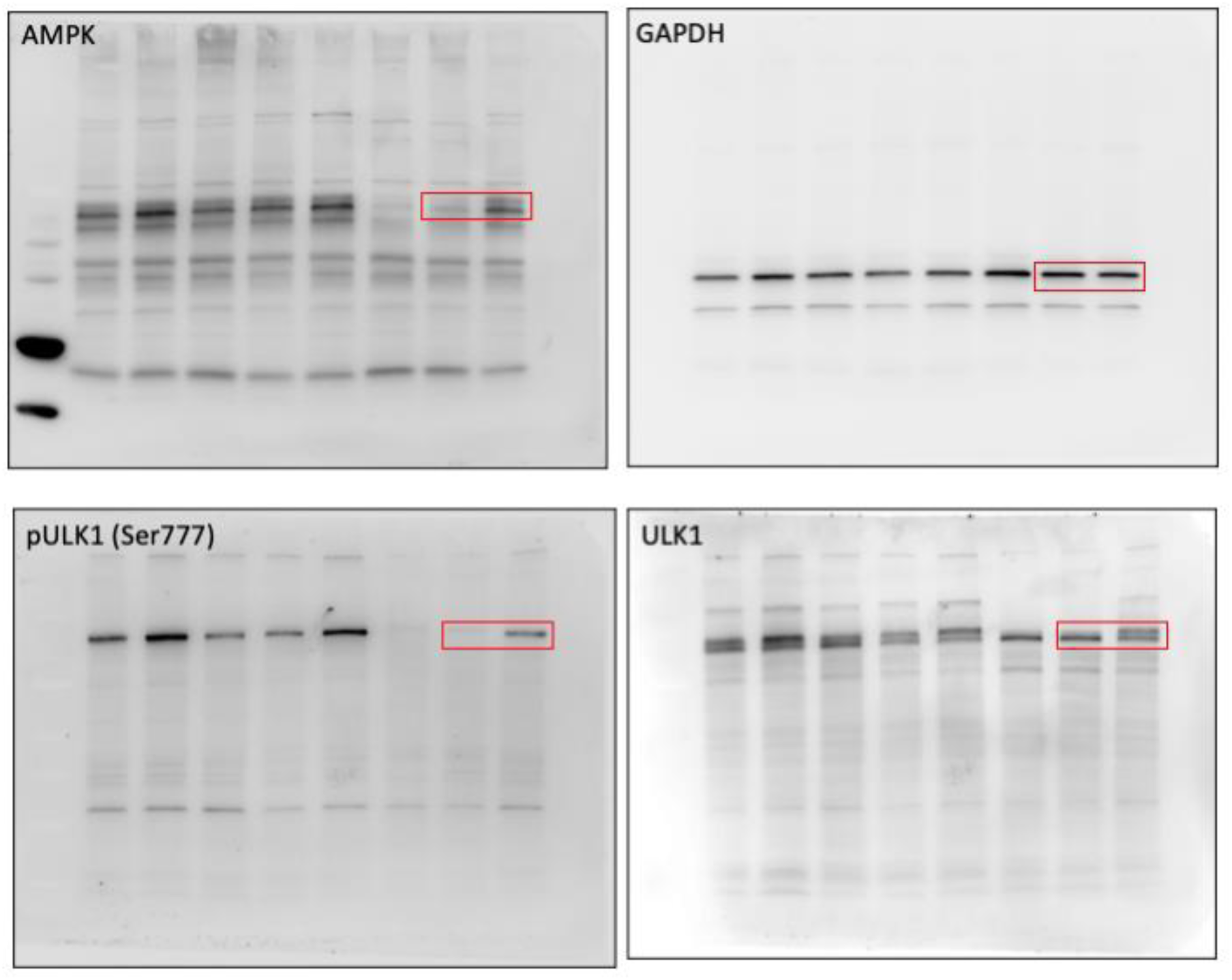
Uncropped western blot image of SCA3 zebrafish treated with SB. Red boxes depict representative images used in Figure 6B.

## References

1. Costa, M.d.C. and H.L. Paulson, Toward understanding Machado-Joseph disease. Progress in neurobiology, 2012. 97(2): p. 239–257.

2. Rüb, U., et al., Clinical features, neurogenetics and neuropathology of the polyglutamine spinocerebellar ataxias type 1, 2, 3, 6 and 7. Prog Neurobiol, 2013. 104: p. 38–66.

3. Privett, B.K. and R.H. Kardon, Spinocerebellar Ataxia with Ophthalmoplegia: 46-y.o. male presenting with progressive esotropia. 2010.

4. Durr, A., et al., Spinocerebellar ataxia 3 and Machado-Joseph disease: clinical, molecular, and neuropathological features. Ann Neurol, 1996. 39(4): p. 490–9.

5. Ranum, L.P., et al., Spinocerebellar ataxia type 1 and Machado-Joseph disease: incidence of CAG expansions among adult-onset ataxia patients from 311 families with dominant, recessive, or sporadic ataxia. American journal of human genetics, 1995. 57(3): p. 603–608.

6. Schols, L., et al., Trinucleotide expansion within the MJD1 gene presents clinically as spinocerebellar ataxia and occurs most frequently in German SCA patients. Hum Mol Genet, 1995. 4(6): p. 1001–5.

7. Bettencourt, C., et al., The (CAG)n tract of Machado-Joseph Disease gene (ATXN3): a comparison between DNA and mRNA in patients and controls. European journal of human genetics : EJHG, 2010. 18(5): p. 621–623.

8. Burt, T., et al., Machado-Joseph disease in east Arnhem Land, Australia: chromosome 14q32.1 expanded repeat confirmed in four families. Neurology, 1996. 46(4): p. 1118–22.

9. Kawaguchi, Y., et al., CAG expansions in a novel gene for Machado-Joseph disease at chromosome 14q32.1. Nat Genet, 1994. 8(3): p. 221–8.

10. Takiyama, Y., et al., The gene for Machado-Joseph disease maps to human chromosome 14q. Nat Genet, 1993. 4(3): p. 300–4.

11. Nascimento-Ferreira, I., et al., Beclin 1 mitigates motor and neuropathological deficits in genetic mouse models of Machado-Joseph disease. Brain, 2013. 136(Pt 7): p. 2173–88.

12. Maciel, P., et al., Correlation between CAG repeat length and clinical features in Machado-Joseph disease. Am J Hum Genet, 1995. 57(1): p. 54–61.

13. Matsumura, R., et al., Relationship of (CAG)nC configuration to repeat instability of the Machado-Joseph disease gene. Hum Genet, 1996. 98(6): p. 643–5.

14. Costa Mdo, C. and H.L. Paulson, Toward understanding Machado-Joseph disease. Prog Neurobiol, 2012. 97(2): p. 239–57.

15. Schmidt, T., et al., An isoform of ataxin-3 accumulates in the nucleus of neuronal cells in affected brain regions of SCA3 patients. Brain Pathol, 1998. 8(4): p. 669–79.

16. Yamada, M., S. Tsuji, and H. Takahashi, Pathology of CAG repeat diseases. Neuropathology, 2000. 20(4): p. 319–25.

17. Goti, D., et al., A mutant ataxin-3 putative-cleavage fragment in brains of Machado-Joseph disease patients and transgenic mice is cytotoxic above a critical concentration. J Neurosci, 2004. 24(45): p. 10266–79.

18. Evert, B.O., et al., Ataxin-3 represses transcription via chromatin binding, interaction with histone deacetylase 3, and histone deacetylation. J Neurosci, 2006. 26(44): p. 11474–86.

19. Evers, M.M., L.J.A. Toonen, and W.M.C. van Roon-Mom, Ataxin-3 protein and RNA toxicity in spinocerebellar ataxia type 3: current insights and emerging therapeutic strategies. Molecular neurobiology, 2014. 49(3): p. 1513–1531.

20. Li, F., et al., Ataxin-3 is a histone-binding protein with two independent transcriptional corepressor activities. J Biol Chem, 2002. 277(47): p. 45004–12.

21. Jung, J. and N. Bonini, CREB-Binding Protein Modulates Repeat Instability in a <em>Drosophila</em> Model for PolyQ Disease. Science, 2007. 315(5820): p. 1857–1859.

22. Chou, A.-H., et al., Polyglutamine-expanded ataxin-3 impairs long-term depression in Purkinje neurons of SCA3 transgenic mouse by inhibiting HAT and impairing histone acetylation. Brain Research, 2014. 1583: p. 220–229.

23. Minamiyama, M., et al., Sodium butyrate ameliorates phenotypic expression in a transgenic mouse model of spinal and bulbar muscular atrophy. Hum Mol Genet, 2004. 13(11): p. 1183–92.

24. Yi, J., et al., Sodium valproate alleviates neurodegeneration in SCA3/MJD via suppressing apoptosis and rescuing the hypoacetylation levels of histone H3 and H4. PloS one, 2013. 8(1): p. e54792–e54792.

25. Chou, A.-H., et al., HDAC inhibitor sodium butyrate reverses transcriptional downregulation and ameliorates ataxic symptoms in a transgenic mouse model of SCA3. Neurobiology of Disease, 2011. 41(2): p. 481–488.

26. Watchon, M., et al., Sodium valproate increases activity of the sirtuin pathway resulting in beneficial effects for spinocerebellar ataxia-3 in vivo. Mol Brain, 2021. 14(1): p. 128.

27. Haggarty, S.J. and L.H. Tsai, Probing the role of HDACs and mechanisms of chromatin-mediated neuroplasticity. Neurobiol Learn Mem, 2011. 96(1): p. 41–52.

28. Liu, H., et al., Butyrate: A Double-Edged Sword for Health? Adv Nutr, 2018. 9(1):p. 21–29.

29. Wu, Y.-L., et al., Treatment with Caffeic Acid and Resveratrol Alleviates Oxidative Stress Induced Neurotoxicity in Cell and Drosophila Models of Spinocerebellar Ataxia Type3. Scientific Reports, 2017. 7(1): p. 11641.

30. Graff, J., et al., An epigenetic blockade of cognitive functions in the neurodegenerating brain. Nature, 2012. 483(7388): p. 222–6.

31. Hockly, E., et al., Suberoylanilide hydroxamic acid, a histone deacetylase inhibitor, ameliorates motor deficits in a mouse model of Huntington’s disease. Proc Natl Acad Sci U S A, 2003. 100(4): p. 2041–6.

32. Kazantsev, A.G. and L.M. Thompson, Therapeutic application of histone deacetylase inhibitors for central nervous system disorders. Nat Rev Drug Discov, 2008. 7(10): p. 854–68.

33. Krainc, D., Clearance of mutant proteins as a therapeutic target in neurodegenerative diseases. Arch Neurol, 2010. 67(4): p. 388–92.

34. Richon, V.M., et al., A class of hybrid polar inducers of transformed cell differentiation inhibits histone deacetylases. Proc Natl Acad Sci U S A, 1998. 95(6): p. 3003–7.

35. Vigushin, D.M., et al., Trichostatin A Is a Histone Deacetylase Inhibitor with Potent Antitumor Activity against Breast Cancer <em>in Vivo</em>. Clinical Cancer Research, 2001. 7(4): p. 971–976.

36. Yeung, F., et al., Modulation of NF-κB-dependent transcription and cell survival by the SIRT1 deacetylase. The EMBO Journal, 2004. 23(12): p. 2369–2380.

37. Göttlicher, M., et al., Valproic acid defines a novel class of HDAC inhibitors inducing differentiation of transformed cells. The EMBO journal, 2001. 20(24):p. 6969–6978.

38. Sealy, L. and R. Chalkley, The effect of sodium butyrate on histone modification. Cell, 1978. 14(1): p. 115–21.

39. Lei, L.F., et al., Safety and efficacy of valproic acid treatment in SCA3/MJD patients. Parkinsonism Relat Disord, 2016. 26: p. 55–61.

40. Canani, R.B., et al., Potential beneficial effects of butyrate in intestinal and extraintestinal diseases. World J Gastroenterol, 2011. 17(12): p. 1519–28.

41. Paganoni, S., et al., Long-term survival of participants in the CENTAUR trial of sodium phenylbutyrate-taurursodiol in amyotrophic lateral sclerosis. Muscle & Nerve, 2021. 63(1): p. 31–39.

42. Ferrante, R.J., et al., Histone deacetylase inhibition by sodium butyrate chemotherapy ameliorates the neurodegenerative phenotype in Huntington’s disease mice. J Neurosci, 2003. 23(28): p. 9418–27.

43. Robinson, K.J., et al., Flow cytometry allows rapid detection of protein aggregates in cellular and zebrafish models of spinocerebellar ataxia 3. Dis Model Mech, 2021. 14(10).

44. Watchon, M., et al., Calpain Inhibition Is Protective in Machado–Joseph Disease Zebrafish Due to Induction of Autophagy. The Journal of Neuroscience, 2017. 37(32): p. 7782–7794.

45. Nascimento-Ferreira, I., et al., Overexpression of the autophagic beclin-1 protein clears mutant ataxin-3 and alleviates Machado-Joseph disease. Brain, 2011. 134(Pt 5): p. 1400–15.

46. Onofre, I., et al., Fibroblasts of Machado Joseph Disease patients reveal autophagy impairment. Sci Rep, 2016. 6: p. 28220.

47. Sittler, A., et al., Deregulation of autophagy in postmortem brains of Machado-Joseph disease patients. Neuropathology, 2018. 38(2): p. 113–124.

48. Mori, F., et al., Autophagy-related proteins (p62, NBR1 and LC3) in intranuclear inclusions in neurodegenerative diseases. Neurosci Lett, 2012. 522(2): p. 134–8.

49. Watchon, M., et al., Autophagy function and benefits of autophagy induction in models of spinocerebellar ataxia type 3. Cells, 2023. 12(6): p. 893.

50. Lee, A., et al., Liver membrane proteome glycosylation changes in mice bearing an extra-hepatic tumor. Mol Cell Proteomics, 2011. 10(9): p. M900538MCP200.

51. Zybailov, B., et al., Statistical analysis of membrane proteome expression changes in Saccharomyces cerevisiae. J Proteome Res, 2006. 5(9): p. 2339–47.

52. Kim, J., et al., AMPK and mTOR regulate autophagy through direct phosphorylation of Ulk1. Nat Cell Biol, 2011. 13(2): p. 132–41.

53. Chou, A.-H., et al., Polyglutamine-expanded ataxin-3 causes cerebellar dysfunction of SCA3 transgenic mice by inducing transcriptional dysregulation. Neurobiology of Disease, 2008. 31(1): p. 89–101.

54. Hockly, E., et al., Suberoylanilide hydroxamic acid, a histone deacetylase inhibitor, ameliorates motor deficits in a mouse model of Huntington’s disease. Proceedings of the National Academy of Sciences, 2003. 100(4): p. 2041–2046.

55. Breuer, P., et al., Nuclear aggregation of polyglutamine-expanded ataxin-3: fragments escape the cytoplasmic quality control. J Biol Chem, 2010. 285(9): p. 6532–7.

56. Qiao, C.M., et al., Sodium butyrate causes alpha-synuclein degradation by an Atg5-dependent and PI3K/Akt/mTOR-related autophagy pathway. Exp Cell Res, 2019: p. 111772.

57. Kakoty, V., et al., Neuroprotective Effects of Trehalose and Sodium Butyrate on Preformed Fibrillar Form of alpha-Synuclein-Induced Rat Model of Parkinson’s Disease. ACS Chem Neurosci, 2021. 12(14): p. 2643–2660.

58. Chen, P.S., et al., Valproate protects dopaminergic neurons in midbrain neuron/glia cultures by stimulating the release of neurotrophic factors from astrocytes. Molecular Psychiatry, 2006. 11(12): p. 1116–1125.

59. Leng, Y. and D.-M. Chuang, Endogenous α-Synuclein Is Induced by Valproic Acid through Histone Deacetylase Inhibition and Participates in Neuroprotection against Glutamate-Induced Excitotoxicity. The Journal of Neuroscience, 2006. 26(28): p. 7502–7512.

60. Vidoni, C., et al., Resveratrol protects neuronal-like cells expressing mutant Huntingtin from dopamine toxicity by rescuing ATG4-mediated autophagosome formation. Neurochem Int, 2018. 117: p. 174–187.

61. Li, X.Z., et al., Therapeutic effects of valproate combined with lithium carbonate on MPTP-induced parkinsonism in mice: possible mediation through enhanced autophagy. Int J Neurosci, 2013. 123(2): p. 73–9.

62. Wang, X., et al., Valproate Attenuates 25-kDa C-Terminal Fragment of TDP-43-Induced Neuronal Toxicity via Suppressing Endoplasmic Reticulum Stress and Activating Autophagy. Int J Biol Sci, 2015. 11(7): p. 752–61.

63. Zhang, Y., et al., Valproic acid protects against MPP(+)-mediated neurotoxicity in SH-SY5Y Cells through autophagy. Neurosci Lett, 2017. 638: p. 60–68.

64. Morselli, E., et al., Spermidine and resveratrol induce autophagy by distinct pathways converging on the acetylproteome. The Journal of cell biology, 2011. 192(4): p. 615–629.

65. Luo, S., et al., Sodium butyrate induces autophagy in colorectal cancer cells through LKB1/AMPK signaling. J Physiol Biochem, 2019. 75(1): p. 53–63.

66. Wang, F., et al., Sodium butyrate inhibits migration and induces AMPK-mTOR pathway-dependent autophagy and ROS-mediated apoptosis via the miR-139-5p/Bmi-1 axis in human bladder cancer cells. FASEB J, 2020. 34(3): p. 4266–4282.

67. Yoo, H.Y., et al., Regulation of Butyrate-Induced Resistance through AMPK Signaling Pathway in Human Colon Cancer Cells. Biomedicines, 2021. 9(11).

68. Zhang, J., et al., Histone deacetylase inhibitors induce autophagy through FOXO1-dependent pathways. Autophagy, 2015. 11(4): p. 629–42.

69. Yun, H., et al., AMP-activated protein kinase mediates the antioxidant effects of resveratrol through regulation of the transcription factor FoxO1. FEBS J, 2014. 281(19): p. 4421–38.

70. Kenyon, C., et al., A C. elegans mutant that lives twice as long as wild type. Nature, 1993. 366(6454): p. 461–464.

71. Gottlieb, S. and G. Ruvkun, daf-2, daf-16 and daf-23: genetically interacting genes controlling Dauer formation in Caenorhabditis elegans. Genetics, 1994. 137(1): p. 107–120.

72. Delpoux, A., et al., FOXO1 constrains activation and regulates senescence in CD8 T cells. Cell Reports, 2021. 34(4): p. 108674.

73. Neri, C., Role and Therapeutic Potential of the Pro-Longevity Factor FOXO and Its Regulators in Neurodegenerative Disease. Front Pharmacol, 2012. 3: p. 15.

74. Gomora-Garcia, J.C., et al., Effect of the Ketone Body, D-beta-Hydroxybutyrate, on Sirtuin2-Mediated Regulation of Mitochondrial Quality Control and the Autophagy-Lysosomal Pathway. Cells, 2023. 12(3).

75. Teixeira-Castro, A., et al., Neuron-specific proteotoxicity of mutant ataxin-3 in C. elegans: rescue by the DAF-16 and HSF-1 pathways. Hum Mol Genet, 2011. 20(15): p. 2996–3009.

76. Vasconcelos-Ferreira, A., et al., ULK overexpression mitigates motor deficits and neuropathology in mouse models of Machado-Joseph disease. Mol Ther, 2022. 30(1): p. 370–387.

77. Ashkenazi, A., et al., Polyglutamine tracts regulate beclin 1-dependent autophagy. Nature, 2017. 545(7652): p. 108-111.

78. Cunha-Santos, J., et al., Caloric restriction blocks neuropathology and motor deficits in Machado-Joseph disease mouse models through SIRT1 pathway. Nature communications, 2016. 7: p. 11445–11445.

79. Lee, J.H., et al., n-Butylidenephthalide Modulates Autophagy to Ameliorate Neuropathological Progress of Spinocerebellar Ataxia Type 3 through mTOR Pathway. Int J Mol Sci, 2021. 22(12).

80. Lin, C.H., et al., Novel Lactulose and Melibiose Targeting Autophagy to Reduce PolyQ Aggregation in Cell Models of Spinocerebellar Ataxia 3. CNS Neurol Disord Drug Targets, 2016. 15(3): p. 351–9.

81. Marcelo, A., et al., Cordycepin activates autophagy through AMPK phosphorylation to reduce abnormalities in Machado-Joseph disease models. Hum Mol Genet, 2019. 28(1): p. 51–63.

82. Ou, Z., et al., Autophagy Promoted the Degradation of Mutant ATXN3 in Neurally Differentiated Spinocerebellar Ataxia-3 Human Induced Pluripotent Stem Cells. Biomed Res Int, 2016. 2016: p. 6701793.

83. Vasconcelos-Ferreira, A., et al., The autophagy-enhancing drug carbamazepine improves neuropathology and motor impairment in mouse models of Machado-Joseph disease. Neuropathol Appl Neurobiol, 2022. 48(1): p. e12763.

84. Wu, J.C., et al., The regulation of N-terminal Huntingtin (Htt552) accumulation by Beclin1. Acta Pharmacol Sin, 2012. 33(6): p. 743–51.

85. Robinson, K.J., et al., A Novel Calpain Inhibitor Compound Has Protective Effects on a Zebrafish Model of Spinocerebellar Ataxia Type 3. Cells, 2021. 10(10).

86. Santana, M.M., et al., Trehalose alleviates the phenotype of Machado-Joseph disease mouse models. J Transl Med, 2020. 18(1): p. 161.

87. Zaltzman, R., et al., Trehalose in Machado-Joseph Disease: Safety, Tolerability, and Efficacy. Cerebellum, 2020. 19(5): p. 672–679.

88. Nandwana, V., et al., The Role of Microbiome in Brain Development and Neurodegenerative Diseases. Molecules, 2022. 27(11).

89. Singh, A., T.M. Dawson, and S. Kulkarni, Neurodegenerative disorders and gut-brain interactions. J Clin Invest, 2021. 131(13).

90. Gamage, H., et al., Machado Joseph disease severity is linked with gut microbiota alterations in transgenic mice. Neurobiol Dis, 2023. 179: p. 106051.

91. Zhang, Y.G., et al., Target Intestinal Microbiota to Alleviate Disease Progression in Amyotrophic Lateral Sclerosis. Clin Ther, 2017. 39(2): p. 322–336.

92. Saute, J.A., et al., A randomized, phase 2 clinical trial of lithium carbonate in Machado-Joseph disease. Mov Disord, 2014. 29(4): p. 568–73.

93. Westerfield, M., The Zebrafish Book : A Guide for the Laboratory Use of Zebrafish. http://zfin.org/zf_info/zfbook/zfbk.html, 2000.

94. Chai, Y., et al., Analysis of the role of heat shock protein (Hsp) molecular chaperones in polyglutamine disease. J Neurosci, 1999. 19(23): p. 10338–47.

95. Cunliffe, V.T., Zebrafish: A Practical Approach. *Edited by* C. NÜSslein-Volhard and R. Dahm. Oxford University Press. *2002. 322 pages. ISBN 0 19 963808 X. Price £40.00 (paperback). ISBN 0 19 963809 8. Price £80.00 (hardback)*. Genetical Research, 2003. 82(1): p. 79–79.

96. Whiten, D.R., et al., Rapid flow cytometric measurement of protein inclusions and nuclear trafficking. Scientific Reports, 2016. 6(1): p. 31138.

97. Lee, A., et al., Rat liver membrane glycoproteome: enrichment by phase partitioning and glycoprotein capture. J Proteome Res, 2009. 8(2): p. 770–81.

98. Perez-Riverol, Y., et al., The PRIDE database and related tools and resources in 2019: improving support for quantification data. Nucleic Acids Res, 2019. 47(D1): p. D442–D450.

